# Intestinal bacteria hijack an evolutionarily conserved epithelial repair signal

**DOI:** 10.64898/2026.08.11.744130

**Authors:** Sam Benson, Priscilla Chin, Sebastian Rogatti, Alessandro Scopelliti, Luigi Zechini, Rosalind Heron, Charlotte Dawson, Anna Davey, Mariska Simpson, Amanda O. Wong, Jair Marques, Alexander von Kriegsheim, David H. Dockrell, Christopher D. Lucas, Jenna Cash, Will Wood, Christopher J. Anderson

## Abstract

Apoptosis in the gut triggers an expansion in the Enterobacteriaceae family of bacteria, causing both prolonged tissue injury and delayed repair. However, the mechanisms linking the Enterobacteriaceae bloom and subsequent deleterious tissue response are relatively unknown. Here, we establish purines as a major component of the apoptotic secretome that are consumed by bacteria. Explicitly, we identify hypoxanthine as a critical metabolite that is taken up and metabolised by both pathogenic and commensal species within the Enterobacteriaceae family. Epithelial cells release hypoxanthine into the extracellular space during early stages of apoptosis via the upregulation of equilibrative nucleoside transporters 1/2 (ENT1/2). Critically, beyond simply linking host and microbe, we delineate a connection between the release of hypoxanthine from the dying cell and the ability of the host to repair damaged epithelial tissue. Hypoxanthine is a potent promoter of epithelial cell repair in both gut and skin across the phylogenetic tree including humans, mice, zebrafish, and fruit flies and promotes similar ATP production and cellular proliferation in eukaryotic and microbial recipients. Thus, the preferential utilisation of hypoxanthine by the Enterobacteriaceae directly competes with the host for a core reparative signal.

## Introduction

Competition for nutrients is constantly raging between mammalian host cells and the microbiota. This competition reaches its zenith in the colon, with a bacterial load of ∼10^11^ bacteria mL^-1^, all occupying their own metabolic and physiological niche (*1*). At homeostasis the population dynamics in the microbiota remains relatively stable. Yet, when apoptosis is induced in response to tissue-damaging insults such as cytotoxic chemotherapies (*2*), inflammatory disorders including Inflammatory Bowel Disease (IBD) (*3*), and physical wounds (*4*), new metabolic niches arise and a common comorbidity is an outgrowth of the Enterobacteriaceae family of bacteria (*5*). The expansion of Enterobacteriaceae is driven in part by death-induced nutrient release (DINNR), a vast metabolite efflux from those cells undergoing apoptosis (*6*). Apoptotic cells have been identified as central hubs for host cell-cell communication by providing ‘find-me’ signals to elicit phagocyte recruitment (*7*, *8*) and, more recently, ‘good-bye’ signals to maintain anti-inflammatory states (*9*); however, the identity of apoptosis-dependent signals and their functional relevance remain comparatively underexplored. The exploitation of apoptosis by the Enterobacteriaceae is both concurrent and causative for lengthening recovery times and preventing tissue repair caused by the initial injury (*10*, *11*). Yet, the mere presence of Enterobacteriaceae family members themselves is not inherently damaging to the host; *Escherichia coli,* for example, is a common commensal and non-pathogenic species within the homeostatic microbiota and can actively protect the host from colonisation of pathogenic species (*12*, *13*). This raises the question: what is it about this unbalanced microbial community, particularly the DINNR-dependent Enterobacteriaceae bloom, that negatively influences tissue repair? Well-established links between a perturbed microbiota and subsequent tissue disease exist, including reduced production of microbially-derived tissue protective metabolites like the short-chain fatty acid butyrate (*14–16*) and the Enterobacteriaceae triggering pro-inflammatory signalling cascades (*17–19*); however, the concept that the Enterobacteriaceae bloom may be competing with the host for injury-resolving factors remains largely untested.

## Results

### Purine salvage drives intestinal E. coli expansion

To begin to investigate the factors that allow for Enterobacteriaceae expansion, we monitored the metabolites released following cytotoxic chemotherapy treatment *in vivo*. For this, wildtype mice that were culture-negative for Enterobacteriaceae were treated with a single dose of doxorubicin with or without oral inoculation with *E. coli* (human commensal strain HS) and compared to naïve untreated controls (Fig. 1A). Doxorubicin-treated mice permitted robust *E. coli* colonisation compared to untreated controls within fresh faecal samples (Fig. 1B) and tissue homogenates (fig. S1A) with similar kinetics as endogenous *E. coli* culture-positive animals (*10*). Semi-targeted metabolomics of the faecal pellets, including a panel of 389 metabolites (Supp table 1), revealed aggregated peak metabolite release occurring 24 hours post-doxorubicin administration (fig. S1B-C). Pathway analysis of metabolite abundance in faecal pellets of doxorubicin-treated mice compared to untreated controls revealed purines to be the most significantly upregulated family of metabolites, followed shortly by pyrimidines (Fig. 1C). This is consistent with previously published mucosal metabolomics data from IBD patients, wherein purine availability is increased in patients with active disease (*20*). Specifically, we observed multiple mono-, di-, and triphosphate nucleotides, including the known apoptosis-dependent ‘find-me’ signal ATP, to be increased in doxorubicin-treated faecal samples compared to naïve controls (Fig. 1D). When comparing mice treated with doxorubicin vs mice treated with doxorubicin and *E. coli*, intestinal *E. coli* colonisation significantly reduced a select group of metabolites (Fig. 1E), including purines (fig. S1D), suggesting their potential consumption by *E. coli*. Notably, the *E. coli* bloom significantly reduced the abundance of core purine intermediates, specifically inosine monophosphate (IMP) and hypoxanthine (Fig. 1F).

**Figure 1.**
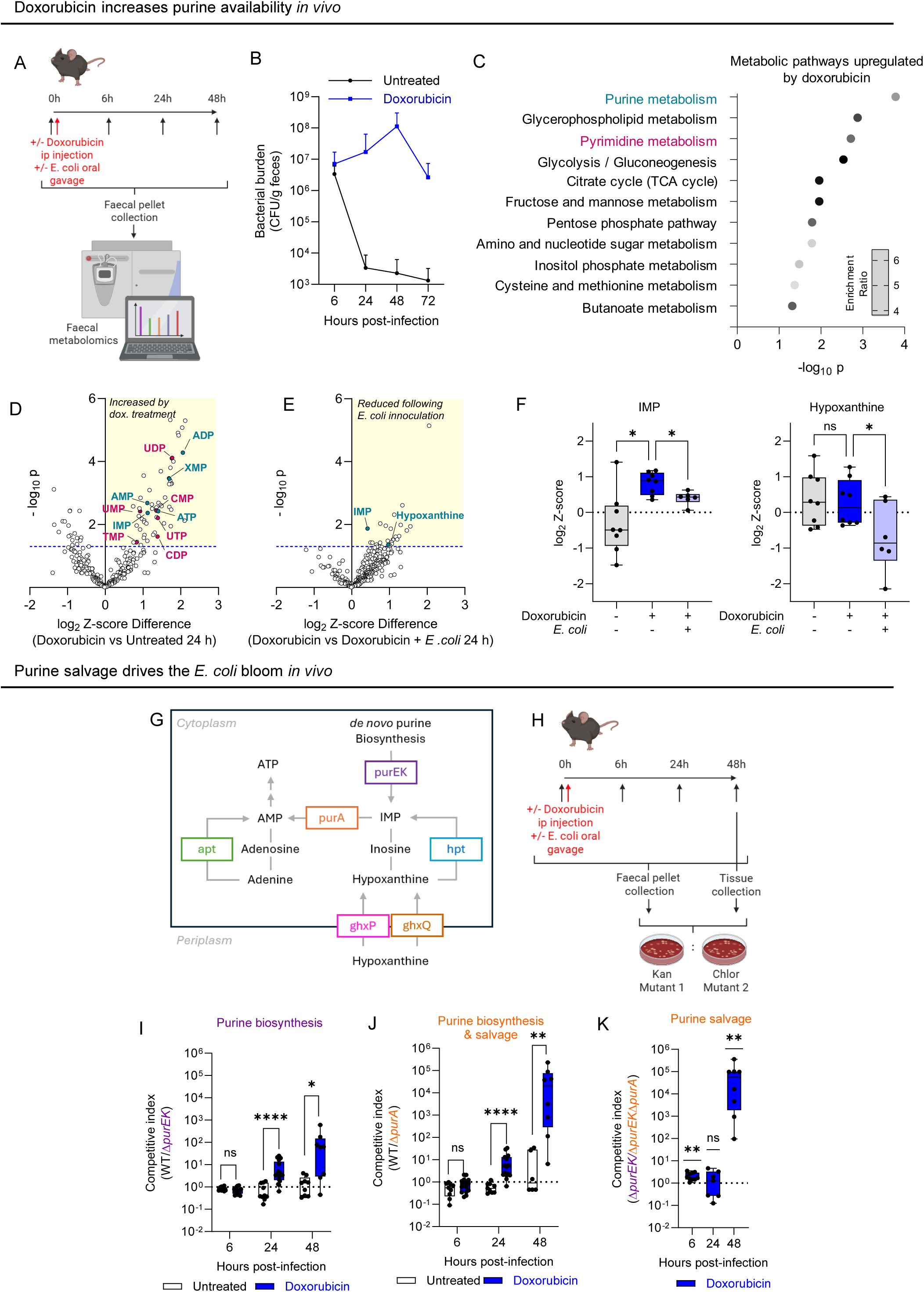
Doxorubicin induced injury increases purine release *in vivo* and salvage by *E. coli*. **A**) Schematic of *in vivo* metabolomics set-up. **B**) Faecal bacterial burden of mice with or without doxorubicin (15 mg kg^-1^). n=10. **C**) Enrichment analysis of statistically upregulated faecal metabolites of mice with or without doxorubicin (15 mg kg^-1^) at 24h post-treatment. **D**) Metabolites released following doxorubicin treatment and **E**) those reduced by the presence of *E. coli*. Purines highlighted in cyan. Pyrimidines highlighted in magenta. Dotted line denotes statistical significance. n ≥ 6, students t tests. **F**) Log_2_-Z-score of faecal IMP & hypoxanthine extracted from D-E. n ≥ 6, one-way Anova. **G**) Schematic of relevant genes and metabolites in purine metabolism within *E. coli.* **H**) Schematic of *in vivo E. coli* competition assay. **I**) Competitive index in faecal pellets between *E. coli* HS WT (SR007) and *ΔpurEK* (PC055). n ≥ 9, multiple Mann-Whitney tests. & **J**) *E. coli* HS WT (SR007) and *ΔpurA* (PC118). n ≥ 6, multiple Mann-Whitney tests. & **K**) *ΔpurEK* (PC055) and *ΔpurEK-purA* (PC120) in doxorubicin treated mice (blue). n ≥ 9, one sample Wilcoxon test.

Purines are essential building blocks for nucleic acid (DNA) synthesis and energy (ATP/GTP) production, with most bacteria (including *E. coli*) capable of both *de novo* biosynthesis and salvage (*21*). Interestingly, IMP, inosine, and hypoxanthine are central components of purine salvage, with IMP acting as a critical branch point for either adenosine monophosphate (AMP) or guanosine monophosphate (GMP) production. Bacterial *de novo* purine biosynthesis requires, among others, PurE and PurK (*22*), to arrive at IMP. The conversion of IMP through to AMP and subsequently ATP requires PurA (*23*). We generated *ΔpurEK* mutant *E. coli* that lacks the ability to create purines *de novo* but can still salvage them, and a *ΔpurA* mutant cannot use purines from *de novo* or salvage routes (Fig. 1G). We inoculated doxorubicin-treated or naïve mice with equal mixtures of wildtype and mutant strains of *E. coli* and monitored bacterial fitness over time (Fig. 1H). In control naïve mice, wildtype and τ1*purEK* (*de novo* purine biosynthesis deficient) were recovered at equal ratios throughout the experiment, indicating that the requirement for *de novo* purine biosynthesis was not generally pertinent to intestinal colonisation. In contrast, in doxorubicin treated mice, wildtype *E. coli* was significantly more fit compared to the purine biosynthesis deficient strain (τ1*purEK*) starting at 24 hours post-inoculation, with an approximate 100-fold advantage observed by 48 hours in faecal pellets (Fig. 1I). Similar fitness values were observed comparing faecal pellets to intestinal tissue homogenates at 24 (fig. S1E-F) and 48 hours (fig. S1G) post-infection. Interestingly, in doxorubicin treated mice, wildtype *E. coli* was a staggering 10,000-fold more fit than the *purA* mutant (*de novo* purine biosynthesis and purine salvage deficient) during peak bloom in faeces (Fig. 1J) and tissue homogenates (fig. S1H), suggesting an integral role for purine salvage.

Given that PurA function is a critical step regardless of if IMP was produced via the purine biosynthesis or salvage pathways, we competed τ1*purEK* with τ1*purEK*τ1*purA* so that neither strain had the ability to synthesise purines *de novo*, allowing for the direct comparison of PurA-mediated salvage to bacterial fitness. The τ1*purEK* ‘control’ strain displayed a similar 10,000-fold fitness advantage over the τ1*purEK*τ1*purA* mutant within the faeces (Fig. 1K) and cecum (fig. S1I) during peak bloom (48 hours), with no τ1*purEK*τ1*purA* recovered from either the ileum or colon. Thus, purine salvage is a critical driver of the *E. coli* bloom *in vivo*, relegating the need for *de* novo purine biosynthesis, and these findings implicate a core group of purine metabolites (IMP, inosine, hypoxanthine) in this process.

### Hypoxanthine recycling fuels E. coli growth

To test whether these core purine metabolites were emanating from apoptotic cells, we turned to an *in vitro* system in which monocultures of human (HCT116) epithelial cells were treated with cytotoxic agents, cell-free supernatants collected, and bacterial growth assessed. We compared the metabolite profiles of supernatants taken directly from dying epithelial cells or taken after *E. coli* growth in apoptotic supernatants to identify which purines are secreted by the dying host cells and potentially consumed by bacteria (Fig. 2A). Intriguingly, only inosine and hypoxanthine were consistently released from dying cells across mouse (CT26) and human (HCT116) host cell lines and distinct apoptotic triggers (doxorubicin or staurosporine) (Fig. 2B, fig. S2A). While levels of inosine and IMP remained constant (fig. S2B), hypoxanthine levels significantly decreased after *E. coli* growth (Fig. 2C), suggesting preferential utilisation of host-derived hypoxanthine. To test if this was a feature specific to this particular commensal strain of *E. coli* or was instead a shared feature of the Enterobacteriaceae family, we performed the same experiment using pathogenic Adherent Invasive *E. coli* (AIEC), the intestinal commensal and systemic pathogen *Klebsiella pneumoniae* (*Klebsiella*), or the foodborne pathogen *Salmonella enterica* serovar Typhimurium (*Salmonella*). Strikingly, hypoxanthine levels were significantly reduced following growth of all species tested (Fig 2D), which displayed similar growth increases in apoptotic supernatants (fig S2C). In contrast, levels of inosine and IMP were unaffected by the growth of these Enterobacteriaceae family members (fig. S2D). These findings suggest that preferential host-derived hypoxanthine utilisation is a fundamental trait of the Enterobacteriaceae, irrespective of strain, species, or pathogenicity.

**Figure 2.**
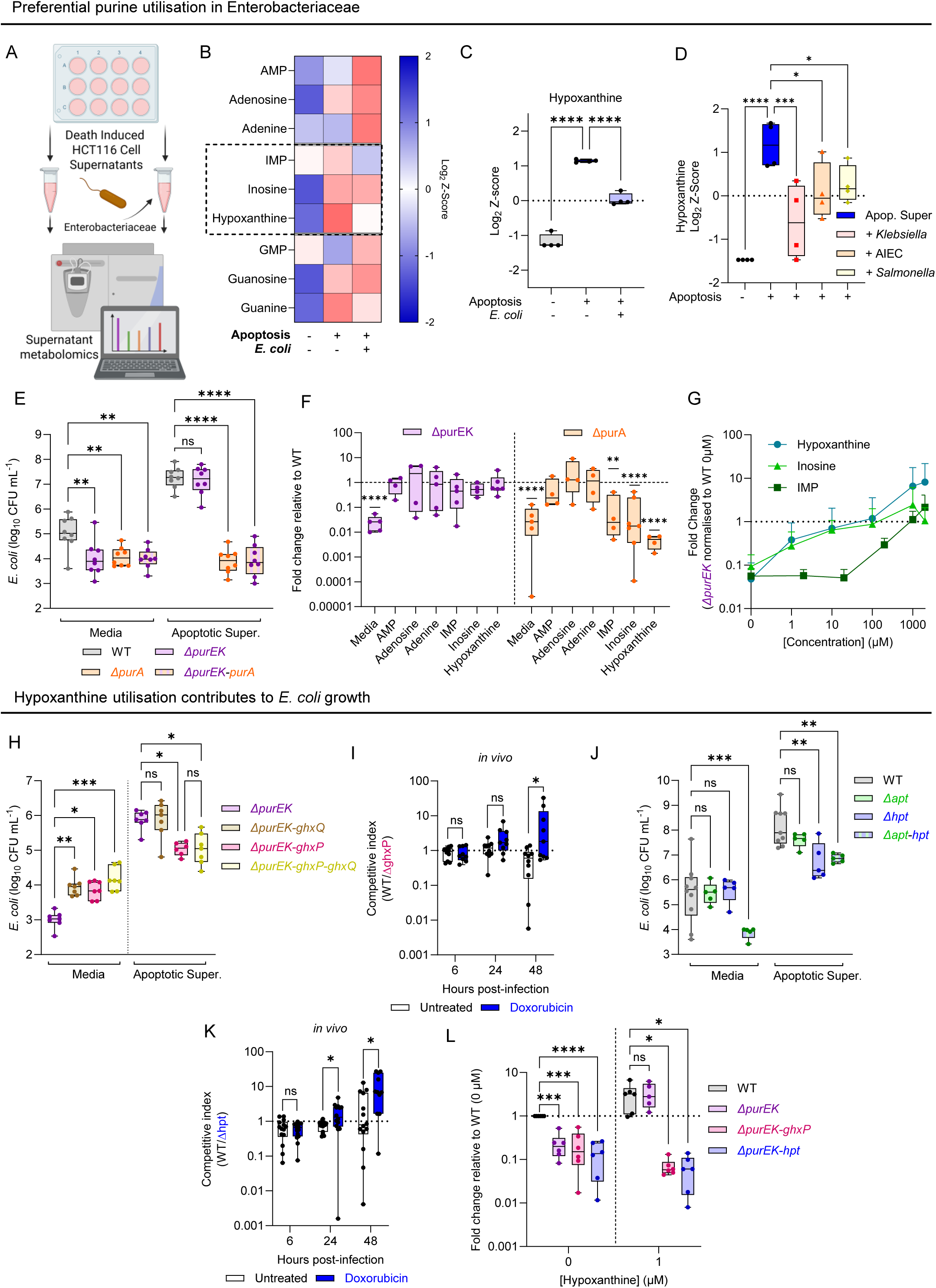
**Enterobacteriaceae preferentially utilise hypoxanthine to promote growth**. **A**) Schematic of *in vitro* metabolomics set-up. **B**) Heatmap displaying purine levels in media controls, supernatants from HCT116 cells treated with Staurosporine (1 μM) for 6h & the same supernatants subsequently inoculated with *E. coli* for 7h. n = 4. **C**) Log_2_-Z-score of hypoxanthine extracted from B. n = 4, one-way Anova. **D**) Log_2_-Z-score of hypoxanthine in apoptotic supernatants inoculated with *K. pneumoniae*, *E. coli* LF82 & *S. typhimarium* SL1344. n=4, one-way Anova. **E**) CFU mL^-1^ of *E. coli* HS WT (grey) vs *ΔpurEK* (CJA117, purple), *ΔpurA* (SJB014, orange) & *ΔpurEK-purA* (SJB017, purple & orange) grown for 7h in media controls or apoptotic supernatants (HCT116 cells, Staurosporine (1 μM), 6h) . n =8, one-way Anova. **F**) Fold change in *ΔpurEK* (CJA117, purple) and *ΔpurA* (SJB014, orange) treated +/- purine derivatives (1 mM) compared to *E. coli* HS WT. n ≥ 4, one sample Wilcoxon test. **G**) Growth of *ΔpurEK* (CJA117) treated with IMP, inosine or hypoxanthine (7h) compared to untreated *E. coli* HS WT. n ≥ 5. **H**) CFU mL^-1^ of *ΔpurEK* (CJA117, purple) vs *ΔpurEK-ghxP* (PC177, pink), *ΔpurEK-ghxQ* (PC196, brown) & *ΔpurEK-ghxP-ghxQ* (PC200, yellow) grown for 7h in media controls or apoptotic supernatants (HCT116 cells, Staurosporine (1 μM), 6h) . n ≥ 6, one-way Anova. **I**) Competitive index between *E. coli* HS WT and *ΔghxP* in mice +/- doxorubicin treatment. n ≥ 9, multiple Mann-Whitney tests. **J**) CFU mL^-1^ of *E. coli* HS WT (grey) vs *Δhpt* (SJB042, blue), *Δapt* (SJB038, green) & *Δhpt-apt* (SJB050, blue & green) grown for 7h in media controls & apoptotic supernatants. (HCT116 cells, Staurosporine (1 μM), 6h). n ≥ 5, one-way Anova. **K**) Competitive index between *E. coli* HS WT and *Δhpt* in mice +/- doxorubicin treatment. n ≥ 11, Multiple Mann-Whitney tests. **L**) Growth of *E. coli* HS WT, *ΔpurEK* (CJA117, purple)*, ΔpurEK-ghxP* (PC177, pink) & *ΔpurEK-hpt* (SJB042, blue) treated +/- hypoxanthine (7h). n ≥ 5, one-way Anova.

To test the functional relevance of hypoxanthine, we undertook an extensive bacterial genetics approach in *E. coli.* Unsurprisingly, in the absence of purine biosynthesis, the *purEK* deficient strain had a significant growth defect in media controls; however, apoptotic supernatants significantly promoted the growth of both the wildtype and *purEK* mutant and restored growth of the *purEK* mutant to wildtype levels (Fig. 2E, fig. S3A). This was true for multiple triggers of apoptosis (staurosporine, doxorubicin), human and mouse colonic epithelial cells, and under aerobic and anaerobic bacterial growth conditions (fig. S3A-B). Importantly, this was generally applicable to purine biosynthesis, as an analogous synthesis-deficient *purMN* mutant (*24*) displayed identical growth phenotypes in media controls and rescue in apoptotic supernatants (fig. S3C). In contrast, purine salvage deficient *purA* mutants were completely insensitive to apoptotic supernatants (Fig. 2E) but could be genetically complemented with either inducible (fig. S3D) or endogenous promoter-mediated expression *in trans* (fig. S3E). These results suggest that whilst appropriate purines are released from the dying mammalian cell to restore a purine biosynthesis deficiency (*ΔpurEK*), the defect caused by blocking bacterial purine salvage (*ΔpurA)* is irredeemable by the metabolites present.

By introducing individual purines to base media one by one at supra-physiological levels (1mM), we aimed to discover which metabolites diverged in their ability to support the growth of *purEK* and *purA* deficient strains. Both strains had an initial defect in media compared to wildtype *E. coli*; however, growth of the purine biosynthesis deficient strain (*ΔpurEK*) could be completely rescued by the introduction of hypoxanthine delineated purines as well as by their adenine counterparts. Conversely, purine salvage deficiency (Δ*purA*) could not be recovered by hypoxanthine derivatives (Fig. 2F). Critically, hypoxanthine and, to a lesser extent, inosine were sufficient to increase both wildtype and *ΔpurEK* growth, while the other purine metabolites either restricted growth (adenine and adenosine) or did not enhance wildtype growth (AMP, IMP) (fig. S3F) at the concentrations tested. Furthermore, the *purEK* mutant displayed a distinct preference for hypoxanthine and inosine compared to IMP, recovering from the initial growth defect at far lower purine concentrations (Fig. 2G). This correlates well with the concentrations of hypoxanthine measured in supernatants (fig. S3G), showing that as little as 1uM of hypoxanthine is sufficient to enable the Enterobacteriaceae bloom.

In parallel, we employed a targeted bacterial genetic approach to identify the key purine(s) that drive death-dependent outgrowth of Enterobacteriaceae. For this, a list of purine transporter candidates were identified from transcriptomic analysis of *E. coli* grown in apoptotic supernatants (*10*) and each transporter was sequentially deleted. Given that the purine biosynthesis genes are some of the highest expressed genes *in vitro*, with the entire 14 gene pathway in the top 25% of all detectable transcripts and 10/14 genes expressed in the top 10% of all detectable transcripts (fig. S4A), transporter mutants were generated in a *ΔpurEK* background to better reveal transporters involved in exogenous purine salvage. No growth defect was seen in the absence of *punC*, *adeP*, and *nupG* that are known to transport adenosine, inosine, guanosine, and adenine (*25–28*); however, a significant growth reduction was observed in the absence of *ghxP* (fig. S4B), a guanine and hypoxanthine transporter (*29*). Interestingly, no additional reduction in growth was observed when *ghxQ*, an additional guanine and hypoxanthine transporter, was deleted, suggesting specificity within this family of purine transporters. Deletion of *ghxP* was sufficient to restrict growth in apoptotic supernatants while deletion of *ghxQ* did not provide an additive growth defect (Fig. 2H). Furthermore, deletion of *ghxP* in an otherwise wildtype strain (with endogenous purine biosynthesis intact) led to a significant fitness defect *in vivo* during peak *E. coli* bloom in doxorubicin-treated mice in both faecal samples (Fig. 2I) and tissue lysates (fig. S4C). Collectively, these findings reveal exquisite specificity for hypoxanthine to permit *E. coli* to fully exploit the apoptotic environment.

Following transport across the outer membrane of *E. coli*, nucleotides (e.g. IMP) must be broken down to their nucleosides and nuclear bases in the periplasm prior to transport across the inner membrane (*30*). Hypoxanthine is then recycled to IMP via hypoxanthine phosphoribosyl transferase (Hpt) and adenine is recycled to AMP via adenine phosphoribosyl transferase (Apt) (Fig. 1G). To test the importance of these enzymes for *E. coli* expansion, we deleted *hpt* and *apt* and assessed bacterial growth. In our *in vitro* system, *Δhpt* exhibits a significant growth defect compared to wildtype in apoptotic supernatants whilst *Δap*t does not, in both a wildtype & purine biosynthesis (*ΔpurEK*) deficient background (Fig. 2J, fig. S4D). Moreover, the combinatorial deletion of *apt* and *hpt* (*Δhpt-apt*) made no additional impact, thus suggesting that adenine salvage is not necessary to enable the full growth phenotype, whilst hypoxanthine salvage defects have a deleterious effect on the ability of *E. coli* to exploit death-induced nutrient release. In agreement, *hpt* deficiency led to a significant fitness reduction *in vivo* in doxorubicin-treated mice in both faeces (Fig. 2K) and tissues with no additional defect seen in the *Δhpt-apt* strain (fig. S4E). Importantly, *ghxP* and *hpt* deficient strains were both insensitive to hypoxanthine-mediated rescue (Fig. 2L) whilst maintaining responsiveness to exogenous IMP and inosine (fig. S3F). Collectively, these data strongly implicate *E. coli* preference for host-derived hypoxanthine and identify both transporter (GhxP) and enzymatic processing (Hpt) for utilisation.

### E. coli depends on pyrimidine biosynthesis for death-dependent growth

There is a well-established relationship between purine and pyrimidine nucleotides, as replicating cells require a balance of the two for nucleic acid, energy, and coenzyme production (*31–33*). Thus, it stood to reason that an increase in purine salvage during apoptosis-driven growth would demand a similar salvage or the *de novo* production of pyrimidines to keep pace. Multiple lines of evidence suggest that *E. coli* must generate newly synthesised pyrimidines, as opposed to relying on host-derived pyrimidines for salvage. First, analysis of metatranscriptomics data from doxorubicin-treated animals revealed a significant upregulation in bacterial transcripts associated with pyrimidine ribonucleotide biosynthesis (fig. S5A), which is in contrast to the downregulation of purine biosynthesis transcripts (*10*). Second, a small number of pyrimidine biosynthesis genes are expressed at higher levels when exposed to apoptotic supernatants (fig. S5B) which is, again, in contrast to purine biosynthesis gene expression (*10*). Third, the levels of the majority of pyrimidines were insensitive to the presence of *E. coli in vivo* (fig. S1D) and *in vitro* (fig. S5C). Finally, the growth of *E. coli* mutants deficient for pyrimidine biosynthesis (*ΔpyrBI* & *ΔpyrF*) was largely insensitive to apoptotic supernatants and strains deficient for genes linked to cytosine uptake and salvage (*codBA*) and uracil uptake and salvage (*uraA*, *upp*) had no growth defect in apoptotic supernatants (fig. S5D).

### Host ENT1/2 exports growth-inducing hypoxanthine during early stages of apoptosis

We next addressed the link between programmed apoptosis and hypoxanthine release. Caspase initiated phosphatidylserine exposure is a typical hallmark of cellular apoptosis (*34*). Host epithelial cells maintained membrane integrity (Draq7-negative) and were largely negative for PS exposure (Annexin V staining) by 6 hours post-death induction, yet both wildtype and *purEK* mutant *E. coli* displayed significant growth increases at this time point (Fig. 3A-C, fig. S6A-B). Despite a lack of PS exposure, these host cells were apoptotic, as activation of the apoptotic caspases 3, 7 & 9 could be readily detected by 6 hours (Fig. 3D-E), the cells eventually exposed PS (Fig. 3A-B, fig. S6A-B), and bacterial growth was reduced using the pan-caspase inhibitor QVD (fig. S6C). To ensure that hypoxanthine release was indeed driven by the apoptotic cascade, we generated a modified line of human intestinal epithelial cells expressing an inducible caspase 9, as caspase 9 is an initiator caspase that links cytochrome c release with caspase 3 activation (*35*, *36*). Induction of caspase 9 was sufficient to drive apoptosis (fig. S6D-E), was accompanied by caspase-dependent growth of wildtype and purine biosynthesis-deficient (*ΔpurEK*) *E. coli* (Fig. 3F) and led to caspase-dependent hypoxanthine release (Fig. 3G). These data demonstrate that caspase 9 activation is sufficient to initiate hypoxanthine release and suggest this is part of the early phases of apoptosis.

**Figure 3.**
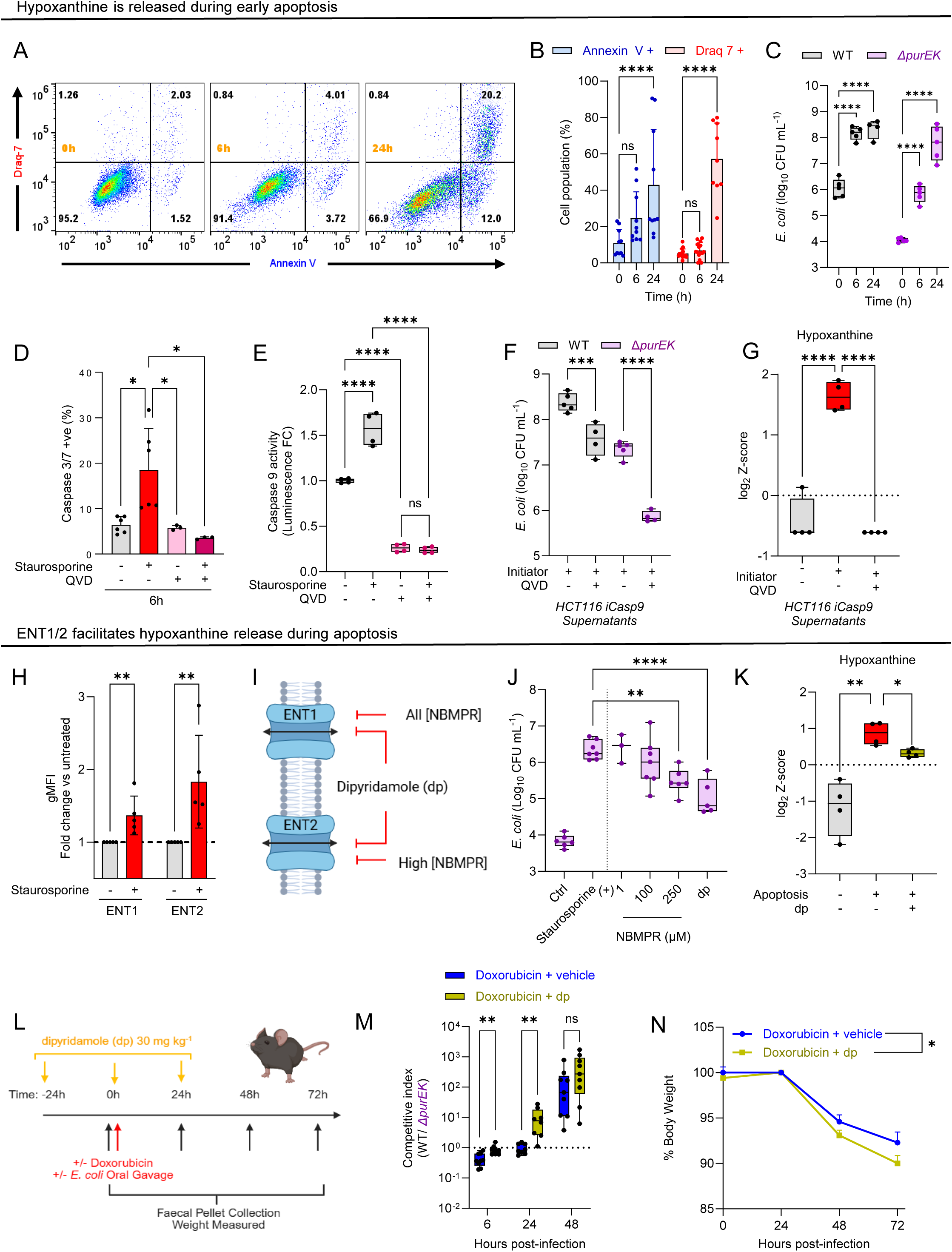
Hypoxanthine release originates in early apoptosis. **A**) Representative flow plots for HCT116 cells treated with staurosporine for 0, 6 or 24h. **B**) levels of Annexin V (blue) and Draq-7 (red) staining in HCT116 cells treated for 0, 6 or 24h with staurosporine. n = 9, one-way Anova. **C**) CFU mL^-1^ of *E. coli* HS WT (grey) vs ΔpurEK (CJA117, purple) grown for 7h in media controls or apoptotic supernatants (HCT116 cells, staurosporine (1 μM), 6/24h). n ≥ 4, one-way Anova. **D**) Caspase 3/7 levels and **E**) caspase 9 levels in HCT116 cells treated with staurosporine for 6h +/- pretreatment with QVD (3 uM). n ≥3, one-way Anova. **F**) CFU mL^-1^ of *E. coli* HS WT (grey) vs ΔpurEK (CJA117, purple) grown for 7h in media controls or apoptotic supernatants (HCT116-*icasp9* cells, initiator AP20187 (2 pM), 6h, +/- QVD (3 uM)). n = 4, one-way Anova. **G**) Relative levels of hypoxanthine in supernatants taken from HCT116 icasp9 cells treated +/- initiator (2 pM) and +/- QVD (3 μM). N = 4, one-way Anova. **H**) ENT1/2 staining in HCT116 cells treated +/- staurosporine for 6h. n = 5, one-way Anova. **I**) Effect of NBMPR and dp on ENT1/2. **J**) CFU mL-1 of *E. coli* HS *ΔpurEK* (CJA117, purple) grown for 7h in media controls or apoptotic supernatants (HCT116 cells, staurosporine (1 uM), 6h, +/- dp (50 uM) +/-NBMPR). n ≥ 3, one-way Anova. **K**) Log_2_-Z-score of hypoxanthine in HCT116 cells treated with staurosporine +/- dipyridamole (50 μM). n=4, one-way Anova. **L**) Treatment plan for mice exposed to dipyridamole during gut injury. **M**) Competitive index between *E. coli* HS WT (SR007) and *ΔpurEK* (PC055) in mice treated with doxorubicin alone (blue) or doxorubicin & dp (30 mg kg-1) (yellow) & **N**) the associated weight loss. n ≥ 9, multiple Mann-Whitney tests.

This left the question of how apoptotic purines are released from the dying cell. Two mammalian equilibrative nucleotide transporters, ENT1 and ENT2 (also known as SLC29A1/2) have been characterized for their ability to transport a range of nucleosides and nuclear bases outside of the cell (*37*). The expression of these transporters on the plasma membrane of epithelial cells is significantly increased during early stages of cell death *in vitro* (Fig. 3H). We chose to inhibit ENT1/2 using two separate drugs. S-(4-Nitrobenzyl)-6-thioinosine (NBMPR), which can block ENT1 alone at lower concentrations and both ENT1/2 at higher concentrations (*38*, *39*), and dipyridamole (dp), which blocks both ENT1/2 at the concentration used (Fig. 3I) (*40*). We observed that NBMPR treatment significantly dampened the growth phenotype of the *de novo* purine biosynthesis deficient *ΔpurEK* strain in a concentration-dependent fashion, with dipyridamole treatment also displaying a significant dampening effect on bacterial growth without influencing host cell death (Fig. 3J, fig. S6F-G). Metabolomic analysis of the dipyridamole-treated apoptotic supernatants demonstrated that blocking ENT1/2 significantly decreased the amount of hypoxanthine released by the dying cell whilst not altering inosine or IMP levels (Fig. 3K, fig. S6G). Dipyridamole-induced bacterial growth reduction could be recovered by the addition of exogenous hypoxanthine and, as a further control, *ΔpurEK-hpt* (hypoxanthine salvage deficient) was insensitive to dipyridamole treatment or exogenous hypoxanthine administration (fig. S6H) thus functionally linking ENT1/2 inhibition to hypoxanthine release.

To test if ENT1/2-mediated purine release influenced *E. coli* outgrowth *in vivo*, we treated mice with dipyridamole via oral gavage prior to and during doxorubicin treatment (Fig. 3L). Dipyridamole treatment increased the requirement for bacterial *de novo* purine biosynthesis, as the fitness advantage for wildtype *E. coli* was significantly increased over the *ΔpurEK* strain even at the early stages of tissue damage (Fig. 3M). However, total bacterial burden levels displayed a notable but not significant overall increase in both faeces and tissue (fig. S6I). Counterintuitively, dipyridamole treatment led to a significant worsening of host disease (Fig. 3N). These findings suggest that blocking ENT1/2-mediated purine release mitigates the ability of *E. coli* to salvage host-derived purines but paradoxically has a detrimental effect on the ability of the host to recover from the initial injury. This posed the question, what purpose might these purines, and specifically hypoxanthine, have on tissue repair and regeneration?

### Hypoxanthine promotes epithelial repair across evolution

To begin to explore the role of hypoxanthine in the host repair process, we employed an epithelial cell scratch assay using human intestinal epithelial cells (Caco-2) and monitored wound closure over 24 hours (Fig. 4A). Addition of exogenous hypoxanthine to the monolayer increased the rate of wound closure and increased cellular ATP production (Fig. 4B-C), suggesting an increase in energy production likely aiding in epithelial cell proliferation. To test if hypoxanthine-mediated repair was consistent across epithelial sites, we utilised an *in vivo* model of acute skin injury, wherein mice received 4 mm dorsal skin excisional wounds and were treated with 30% Pluronic hydrogel containing either hypoxanthine or vehicle (Fig. 4D)(*41*). Topical hypoxanthine administration significantly increased wound re-epithelialisation, with a substantial increase in epithelial tongue length of over 75% compared to vehicle (Fig. 4E) and a corresponding increase in keratinocyte proliferation (Fig. 4F). These data support the notion that hypoxanthine is a critical repair signal for mammalian epithelial cells of both skin and gut origin.

**Figure 4.**
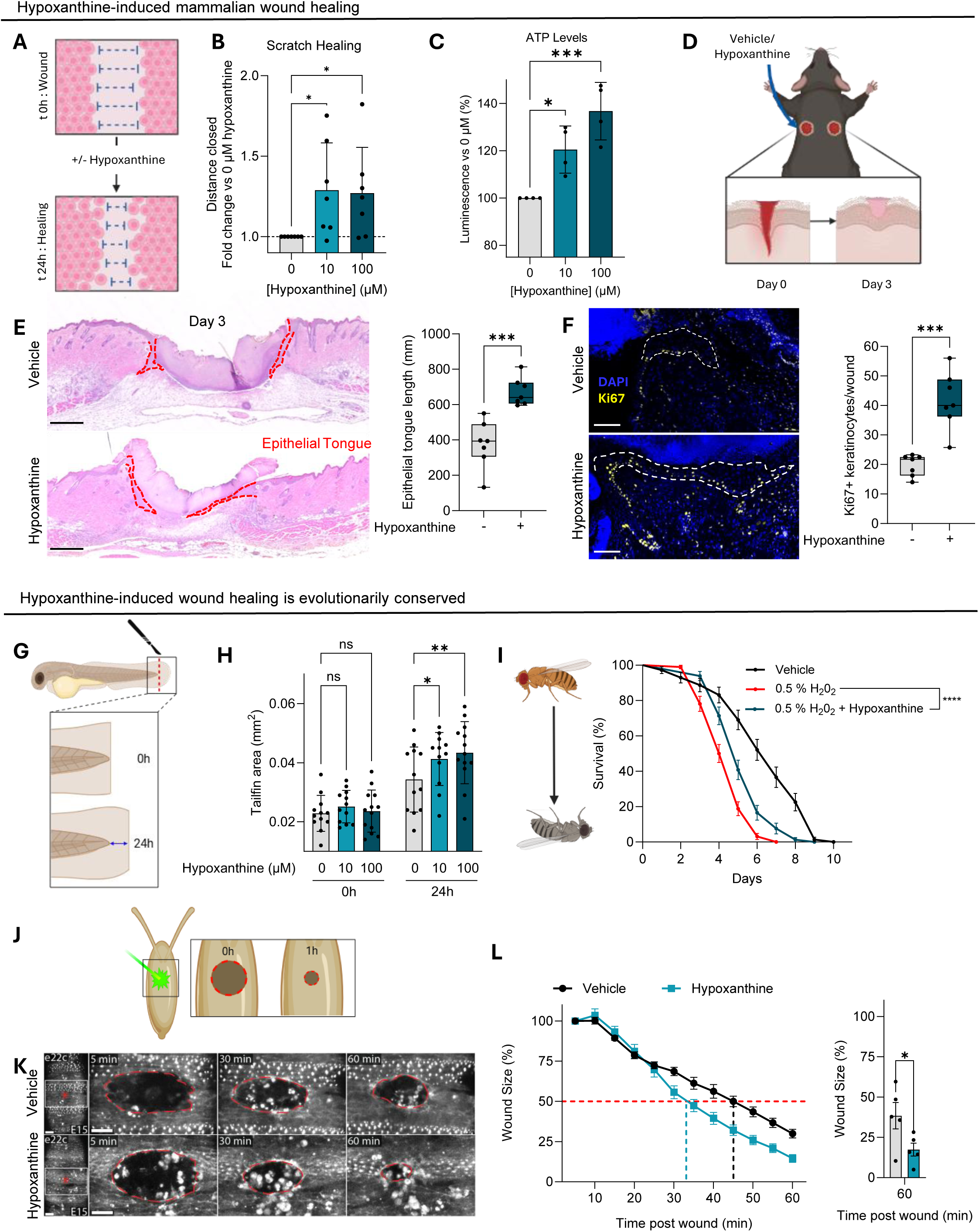
Hypoxanthine is beneficial for wound healing. **A**) Schematic of the scratch assay model of epithelial repair and proliferation using human derived Caco2 cells. **B**) Fold change in scratch closure after 48h when treated with hypoxanthine vs untreated cells. n = 7, one-way Anova. **C**) Fold change in intracellular ATP levels in Caco2 cells exposed to increasing concentrations of hypoxanthine. n = 3, one-way Anova. **D**) Mouse acute skin wound model; 4 mm full skin excisions are carried out to the dorsal flank, and the wound healing process is assessed at 3 dpw. **E**) H&E-stained wound mid-sections at 3 dpw with epithelial tongues highlighted in red. Scale bar 400 μm. Quantification of epithelial tongue length +/- hypoxanthine. n = 7, t-test. **F**) Representative images of Ki67+ keratinocytes within epithelial tongues in vehicle and hypoxanthine-treated wounds 3 dpw. Scale bar 100 μm. Number of Ki67+ keratinocytes per wound. n = 7, t-test. **G**) Zebrafish tail fin regeneration model; tail fin is resected distal to the notochord, recovery monitored over 24 h. **H**) Tailfin area at 0 and 24h post resection. n = 12, two-way Anova. **I**) Survival curve of adult *Drosophila* treated +/- 0.5% H_2_O_2_, +/- hypoxanthine (100 μM). n >71, Kaplan-Meier test. **J**) *Drosophila* embryolaser wound model; injury induced with 355 nm laser, Wound closure monitored over 1 h **K**) Representative images of wound closure, highlighted in red, over 1h +/- hypoxanthine (10 μM). Scale bar: 20 μm. **L**) Percentage wound closure +/- hypoxanthine (10 μM). Dashed lines indicate 50% of the initial wound size (red) and the time to reach this for vehicle (black) and hypoxanthine (teal) treated embryos. n = 5, unpaired t-test.

The ability to salvage purines remains evolutionarily conserved across species (*42*), thus we hypothesised that the reparative properties of hypoxanthine would be similarly conserved. To test this, we first utilised a zebrafish (*Danio rerio*) larvae model of tailfin regeneration. For this, the tailfin is resected distal to the notochord and will regenerate over time (Fig. 4G) (*43*, *44*). We chose this model in part due to the dependency on re-epithelialisation and cellular proliferation for successful repair (*45*, *46*). Tailfin regeneration occurs over time, with detectable regrowth occurring within 24 hours post-resection (fig. S7A) and, in agreement with the mammalian data, hypoxanthine significantly increased the rate of tailfin regrowth (Fig. 4H). To test if hypoxanthine mediates intestinal barrier repair beyond mammalian systems, we turned to a fruit fly (*Drosophila melanogaster*) model of intestinal damage. Feeding adult flies hydrogen peroxide (H_2_O_2_) induces intestinal epithelial damage and subsequent proliferation during injury repair (*47*). Simultaneous feeding with hypoxanthine significantly protected flies from H_2_O_2_-induced death (Fig. 4I) across a range of disease severities (fig. S7B).

To this point, all *in vitro* and *in vivo* models tested are at least in part dependent on cellular proliferation for repair. To determine if hypoxanthine-induced repair was restricted to increased proliferation, we used laser wounding of the *Drosophila* embryo (Fig. 4J), as wound closure in this model is independent of cellular proliferation but instead requires ATP-dependent contraction of an actin-myosin cable around the wound site (*48*). Injection of hypoxanthine into the vitelline space immediately prior to laser wounding promoted significantly faster wound closure over time compared to a vehicle control, with 50% wound closure occurring in >35% less time (Fig. 4 K-L, Supp. Movie 1). Thus, from an initial discovery that inhibiting hypoxanthine release increases damage to the intestinal tract, these findings identify hypoxanthine as a broadly conserved driver of epithelial cell repair.

## Discussion

The release of purines has a long history in the study of apoptosis, but primarily in the form of ATP, a well-known and well-studied ‘find-me’ signal during efferocytosis (*7*). Other purines have a more limited connection with cell death. Hypoxanthine and inosine have been explored as early biomarkers of cell death in myocardial ischemia, concentrations of which are seen to be elevated within minutes of injury, long before more common markers (*49*). In this context it is suggested that mitochondrial degradation accelerates the breakdown of ATP to its catabolic products, a process that may also be occurring in our system. Inosine has also been highlighted in the secretome of apoptotic brown adipocytes for its role in increasing energy expenditure in surrounding adipocytes and the browning of white adipocytes (*50*). But in this scenario, inosine influences cAMP-kinase activity rather than being directly incorporated into ATP. Others have also identified hypoxanthine in supernatants from apoptotic cells but have focused exclusively on upstream metabolites (ATP, AMP, Adenosine, IMP) and their interactions with the immune system (*9*, *51*). Interkingdom transfer of hypoxanthine produced by genetically engineered *E. coli* in the gut to host epithelial cells has been demonstrated to be beneficial for tissue repair within a mouse colitis model, and specifically via an increase in ATP generation, however this beneficial effect is dependent on the complete lack of commensal Enterobacteriaceae (*52–54*). Hypoxanthine-induced wound healing may not be exclusively due to ATP generation, with its degradation previously reported to promote ROS generation leading to faster healing rates (*55*). The ability of almost all forms of life to salvage hypoxanthine for energy conservation is a fundamental metabolic process (*42*). Therefore, almost all cells can theoretically gain some benefit from exogenous hypoxanthine.

Preferential incorporation of death-derived purines by the Enterobacteriaceae emphasises the opportunistic metabolic adaptability inherent to this family of bacteria. By utilising hypoxanthine, the Enterobacteriaceae avoid the energy intensive *de novo* purine biosynthesis pathway, saving 6x ATP alongside other metabolic intermediates inherent in making ATP from scratch (*56*). Such energy efficiency and conservation are, presumably, a critical component of the rapid expansion that occurs *in vivo* during the Enterobacteriaceae bloom in the face of microbial competitors. Although we provide data that purine salvage promotes growth *in vitro* in either aerobic or anaerobic growth conditions, it is less clear how this energy efficiency programme intertwines with the shift to aerobic respiration previously reported *in vivo* (*57*, *58*). Furthermore, it remains unclear if the use of these scavenged purines is the direct driver of bacterial proliferation or a beneficial addition to growth that is initiated by alternative growth substrates. Interestingly, the presence of hypoxanthine appears to be both necessary and sufficient to promote bacterial growth within the context of a glucose-based medium *in vitro*. Regardless, *E. coli* is certainly taking full advantage of the glut of raw materials for purine salvage provided by the dying cell.

The data presented here advance the concept that the Enterobacteriaceae bloom specifically exploits a key apoptosis-derived epithelial repair signal for effective colonisation and outgrowth. In the constant arms race between host and microbe, targeting host factors that play critical roles in tissue homeostasis or tissue repair affords a strategic advantage for the microbial community in that these signals must be produced and present for the host to thrive. Our data suggest that the pilfering of hypoxanthine by the Enterobacteriaceae effectively removes a prominent tissue repair signal and could help explain Enterobacteriaceae bloom-dependent delays in the repair process (*10*, *57*). It remains to be seen if the impact of hypoxanthine is restricted to the epithelial cell lineage or if similar reparative programs would be activated in additional cell types involved in tissue repair and regeneration, such as phagocytic cells. Critically, the reparative nature of hypoxanthine is neither tissue nor species specific, as hypoxanthine improves epithelial repair in the gut and skin across mammals, zebrafish, and fly.

Thus, hypoxanthine is a critical metabolite that is released during early stages of apoptosis, is integral for driving proliferation-dependent and independent wound repair across multiple species spanning the phylogenetic tree, and is preferentially sequestered and utilised by the Enterobacteriaceae to adapt to the host intestinal tract limiting availability for reparative programmes.

## Acknowledgements

We would like to acknowledge the IGC MS facility, the BVS aquatics and rodent staff, and the IRR flow cytometry facility for their technical assistance. We thank Brian McHugh for his support and members of the Edinburgh Cell Death Collective for their insightful comments and suggestions. All schematics were created using BioRender.com. For the purpose of open access, the authors have applied a Creative Commons Attribution (CC BY) license to any Author Accepted Manuscript version arising from this submission.

## Funding

Wellcome Trust Career Development Award 225923/Z/22/Z (CJA) supports SJB, PC, SR, and MS.

Ker Memorial PhD Studentship supports AD.

Wellcome Trust Investigator Award 22460/Z/21/Z (WW) supports LZ and RH.

MRC Programme Grant MR/W019264/1 (WW) supports AS.

MRC Project Grant UKRI2372 (JC) supports CD.

Wellcome Trust (Multiuser Equipment 208402/Z/17) (AvK) supports JM.

## Author Contributions

Conceptualization: SB, CJA. Data Curation: SB.

Formal Analysis: SB, PC, SR, AS, LZ, CD, AD, RH, CD, JC.

Funding acquisition: CJA.

Investigation: SB, CJA, PC, SR, AS, LZ, CD, AD, RH, JM, JC.

Methodology: SB, CJA, PC, AS, LZ, RH, AD, MS, JM, CL, DD, AvK, JC, WW.

Project administration: SB, CJA. Resources: AW.

Supervision: SB, CJA, CL, WW, AvK, JC, DD. Visualization: SB, CJA.

Writing – original draft: SB, CJA, Writing – review & editing: All authors.

## Data, code and materials availability

All data needed to evaluate the conclusions in the paper are present in the paper or the supplementary materials. Metabolome data are available in the MetaboLights database under accession number REQ20260707221380

## Supplementary Materials

Materials & Methods

Figs. S1-S6

Tables S1-S4

References 59-64 Movies S1

## Materials & Methods

### Animal work

All animal work was approved by the University of Edinburgh Animal Welfare and Ethical Review Body (AWERB) and the United Kingdom Home Office (license PP8738752 and PP2067887). C57BL/6 mice were purchased from Charles River and allowed to acclimatise to local animal facilities for a minimum of 1 week prior to experimentation. Mice were confirmed culture-negative for Enterobacteriaceae by plating fresh faecal samples on MacConkey agar without antibiotics prior to inclusion in experiments outlined below. All animal experiments were performed across a minimum of 2 independent cohorts.

#### In vivo mouse model of death-induced Enterobacteriaceae bloom

Doxorubicin was given as a single intraperitoneal injection at 15 mg kg^-1^ of mouse body weight while vehicle control (water) was given at similar volumes (approximately 300 μl volumes of water or 1 mg ml^−1^ doxorubicin solution). Mice were oral gavaged with either *E. coli* HS (1e^8^ in 100 μL PBS) or vehicle control. Mice were weighed daily, including prior to receiving treatment, and fresh faecal samples were taken at indicated times. Tissues were collected at indicated times after treatment. For dipyridamole experiments, dipyridamole (30 mg kg^-1^, 1% carboxymethyl cellulose) or vehicle (1% carboxymethyl cellulose) was administered by oral gavage at day −1, 0 and 1 of the above protocol.

#### In vivo mouse competition assays

Mice were infected via oral gavage with 1e8 CFU of each strain of *E. coli* HS with or without doxorubicin treatment (as described above). Chloramphenicol or kanamycin resistance was achieved by unresolved insertions into the *lacZ* locus as previously described, with the two antibiotic resistance profiles displaying equal fitness (*10*). The ratio of strains in the input was confirmed by plating the infective dose on MacConkey agar plates containing chloramphenicol (final concentration 10mg ml^-1^) and MacConkey agar plates containing kanamycin (final concentration 50mg per ml). Each strain was confirmed to only grow on the expected antibiotic. Dilutions were plated onto MacConkey agar plates containing chloramphenicol (final concentration 10mg ml^-1^) and MacConkey agar plates containing kanamycin (final concentration 50mg ml^-1^) to calculate the ratio of the two strains (output). Competitive indices were calculated as [Output] / [Input] for each sample (faeces, tissues). Given that each exogenously introduced strain was deficient for *lacZ*, and is a non-lactose fermenter, any lactose fermenting colonies were not counted and were viewed as ‘‘passenger’’ endogenous Enterobacteriaceae, though these events were exceptionally rare.

#### Mouse acute wound model

Male mice (7-9 weeks old) were randomly assigned a treatment group and anaesthetized with isoflurane (Zoetis, Leatherhead, UK) by inhalation. Buprenorphine analgesia (0.05 mg/kg, s.c, Vetergesic, Amsterdam) was provided immediately prior to wounding and dorsal hair was removed using a Wahl trimmer. Four full-thickness excisional wounds were made to the shaved dorsal skin using sterile, single use 4 mm punch biopsy tools (Selles Medical, Hull, UK). Vehicle (PBS) or hypoxanthine were delivered topically by pipette into the wound cavity immediately after wounding (40 μl in a 30% Pluronic F-127 gel; liquid at 4 °C, but solidifies at room temperature; Sigma Aldrich). Pluronic hydrogel which was selected for its widespread use as a biocompatible, inert carrier in wound healing studies and its thermosensitive properties that facilitate easy application and retention (*59–61*). Mice were housed with their previous cage mates in a 28 °C warm box (Scanbur, Denmark) overnight following wounding, with paper towels used as bedding to avoid sawdust entering the open wounds. Dome home entrances were enlarged to prevent animals scraping their dorsal skin wounds. Animals were moved into clean conventional cages at 22-24 °C the following morning.

Acute wounds were harvested on days 3 post-wounding. Wounds were fixed in 4% PFA (Sigma Aldrich) overnight at 4 °C on a rocker or at room temperature for 4 h, washed 3 x 5 min in PBS and transferred to 70% ethanol and then embedded in paraffin. Day 3 acute wounds were bisected using a scalpel, embedded in paraffin and sections (10 µm) taken from the middle of the wound.

#### Histology

FFPE sections were deparaffinised and rehydrated in: xylene (2 x 3 min), 100% ethanol (2 x 3 min), 90% ethanol (3 min) and 70% ethanol (3 min). Sections were placed in running tap water before being submerged in haematoxylin (Sigma-Aldrich, Dorset, UK) for 5 s. Excess haematoxylin was rinsed off in tap water twice. Slides were then briefly submerged in acid alcohol (300 ml dH20, 700 ml 100% ethanol, 3 ml concentrated HCL; Sigma-Aldrich), followed by tap water wash. Sections were submerged in eosin (Sigma-Aldrich) for 30 s to provide a counterstain to the haematoxylin. After tap water washes (x 3), sections were dehydrated for mounting in sequential baths of 70%, 90%, 100% ethanol and xylene (x 2) for 30 seconds each. DPX mountant (Sigma-Aldrich) was applied to the sections, followed by a coverslip (25 x 60 mm, 0.13 mm thick; ThermoFisher Scientific, UK). Slides were scanned using an Axioscan Z1 Slide Scanner (Zeiss).

#### Ki67 staining

Opal immunofluorescence staining was used to visualise ki67 in murine wounds. After deparaffinisation, tissue sections were submerged in AR9 Tris EDTA antigen retrieval at 90 C for 15 minutes (Vector laboratories, #H-3301). Slides were then rinsed in TBS-T and tissues were circled using on ImmEdge pen (Vector Labs, #H-4000). Sections were incubated with Ki67 primary antibody (Abcam #ab15580) (0.2 μg mL-1) in a humidified chamber at room temperature for 1 hour. Slides were then washed with TBS-T (3 x 2 min) before being incubated with Opal anti-mouse Polymer HRP for 10 minutes at room temperature (#ARH1001EA), before undergoing another cycle of TBS-T washes. Tissues were then incubated with Opal 570 fluorophore diluted in Amplification Diluent for 10 minutes at room temperature before being counterstained with with Spectral DAPI (Akoya, #SKU-FP1490). Sections were mounted with prolong daimond antifade mountant (Invitrogen, #P36970). Images were taken using the PhenoImager HT (Akoya Biosciences)

#### Epithelial tongue measurements

Epithelial tongues, the portion of epidermis migrating underneath the scab, were measured on H&E-stained wound mid-sections in QuPath using polygon tool. Mean lengths were averaged per wound and per mouse.

#### Zebrafish breeding and maintenance

Breeding/maintenance of adult fish was carried out in accordance with the UK Home Office Animal (Scientific Procedures) Act (1986). 15 adult fish were kept in a tank at 5 fish/litre and supplied with running, filtered water, and fed daily. Adult fish were housed in a 14/10 light/dark cycle at 28.5 degrees Celsius. Fish were bred a maximum of once per week by placing all male and female fish into a mating tank the afternoon before the eggs were required to be collected. The following morning, adult fish were returned to their usual system tank and the eggs laid were collected and stored in the incubator at 28.5 °C in a petri dish filled with 0.5 mg mL^-1^ methylthioninium chloride (methylene blue). Embryos were cleaned and had methylene blue replaced daily.

#### Tailfin model of epithelial injury

Embryos were anaesthetized with 2.5 mL of 40 μg mL^-1^ of MS-222 in a petri dish of methylene blue prior to wounding. Wounding was performed on a glass slide under the microscope using a scalpel to transect a portion of the tailfin at a distance of approximately 1μm distal to the notochord. Embryos were housed in a singular well of a 48 well plate for the duration of the experiment containing methylene blue or appropriate treatment, as indicated in figure legends. Methylene blue was refreshed at least once every 24 h. Images were taken zoomed into the tailfin at 10 x magnification on the Invitrogen™ EVOS™ FL Auto 2 Imaging System using Auto 2 software. Images were analysed using Fiji (ImageJ). The scale was set using an image of a ruler taken at 10x magnification. A line was drawn from the end of the notochord in a straight line 0.1 mm towards the embryo’s head. A new line was then drawn across the 0.1mm end horizontally. The tailfin outline distal to this line was traced, encompassing the wounded/regrowing fin. A second line was then drawn using the pentagon tool to transect the fish’s body horizontally at the very edge of the notochord. The remaining wounded/regrowing tailfin was traced, and this area was measured and recorded in mm^2.^.

#### Drosophila H_2_O_2_ injury

Newly eclosed flies were reared in groups of no more than 30 individuals per vial, with an approximate male-to-female ratio of 1:4. Flies were maintained in temperature and humidity-controlled incubators at 29 °C under a 12 h:12 h light–dark cycle on standard cornmeal-based medium. Culture medium was replaced every 2 days. Mated flies aged 6–9 days were randomly selected and transferred to empty vials containing a glass microfiber filter (Whatman, 1821-021) soaked with 5% sucrose (Fisher, S/8600/53) solution supplemented with graded concentrations of H₂O₂ (0.5%, 1.5%, or 3%, Sigma, 386790), either alone or in combination with 100 μM hypoxanthine. The control solution consisted of 5% sucrose and 2% DMSO (Vehicle, Sigma, D2438). Fresh treatment solutions were provided daily, and mortality was scored manually every 24 h. Only female flies were included in the survival analysis. Dead flies were removed from the vials at each scoring time point, and deceased males were not replaced during the experiment.

#### Drosophila laser wound closure

For labelling of the embryonic epithelium to track the effect of exogenous hypoxanthine on wound closure, e22cGAL4 was used (e22c-gal4) to drive UAS-mCherry-Moesin (*62*).. All fly lines used as part of this work were derived from those ordered through the Bloomington Drosophila Stock Centre (University of Indiana (NIH P40OD018537)). Adult flies were left at 20 °C in an overnight laying cage with fresh yeast paste to promote oviposition and the resulting embryos were collected in cell strainers (Falcon). The embryos were then dechorionated in bleach (Jangro) for 90 s and washed six times with distilled water, before being developmentally staged based on gut. Stage 14 embryos were manually selected and mounted ventral side up on a glass slide with double-sided sticky tape, with a droplet of VOLTALEF oil (VWR Chemicals) added before dehydrating in a sealed box with silica beads for ∼15-30 min at 25 °C as described previously (*63*). Hypoxanthine (10μM) was injected into the perivitelline space at the anterior of the embryo using a FemtoJet 4i Microinjector (Eppendorf) attached to a FemtoJet Injectman 4 (Eppendorf). More VOLTALEF oil was added to each embryo, a No 1.0 coverslip (SLS) was sealed on top and imaging undertaken immediately. Confocal imaging was performed on a Zeiss LSM880 laser scanning confocal microscope equipped with a 40x/1.3 oil immersion objective. The acquisition software used was Zen Blue (Zeiss), with images processed and analysed in FIJI (NIH). The slide was inverted for imaging of the ventral epithelium of the embryo. Epithelial wounds were generated using laser ablation (355 nm pulsed DPSS laser, RAPP OptoElectronic). To monitor wound closure, 10 μm deep Z-stacks were acquired every minute, and the wound area measured at set intervals (5 frames = 5 min) using the Fiji Freehand selections tool. To measure energy generation, embryos expressing ubi-AT1.03NL were excited with a 405 nm laser and 5 μm deep Z-stacks were acquired every 90 s. Emission was collected at 471-489 nm (mse-CFP) and 524-551 nm (cpVenus-FRET). The mse-CFP/cpVenus-FRET ratio was calculated, and fold change relative to the pre-wound image was plotted.

### Metabolomics

*Ex vivo* faecal: Pellets were collected at 0, 24 and 48 h after treatment. Pellets were manually dissociated in extraction buffer (MeCN:MeOH:H_2_O, 5:3:2), 1:20 Weight/Volume. Samples were sonicated for 15 mins prior to centrifugation (13,000G, 10 mins). The supernatants were filtered through a 0.22 μm syringe filter, collected and analysed directly.

*In vitro* supernatant: Supernatants were collected as detailed below. 50 μL of supernatant was diluted 1:20 into the extraction buffer (MeCN:MeOH:H_2_O, 5:3:2). Samples were filtered through a 0.22 μm syringe filter, collected and analysed directly.

Hypoxanthine Quantification: Samples were doped with 10 μM of ^13^C_5_-hypoxanthine as an internal standard and extracted as above. Media containing hypoxanthine (0.5 – 1000 μM) doped and extracted as above.

Data Generation: Metabolites were analysed on a Dionex Ultimate 3000 UHPLC (Thermo Fisher Scientific) coupled to a Q Exactive Hybrid Orbitrap MS (Thermo Fisher Scientific). Hydrophilic interaction liquid chromatography (HILIC) using a ZIC-pHILIC analytical column (2.1 × 150 mm, SeQuant-MerckMillipore) coupled with a guard column (2.1 × 20 mm) was used for chromatographic separation of metabolites. A gradient using mobile phase A 20 mM ammonium carbonate, 0.05 % (v/v) ammonium hydroxide, and B (acetonitrile) was used. The gradient was set from 95 % to 5 % B over 20 min and reequilibrated to initial conditions for 7 min. The flow rate was 200 ml min^-1^, and the temperature was at 45 °C. The injection volume was 5 μl. MS data was acquired in positive/negative polarity switch mode in the m/z range of 70–900 Da, with a resolving power of 70,000 (FWHM).

Analysis: MS peak integration was performed on Skyline to produce relative values for the targeted metabolites. Log_2_-Z-scores were calculated from ½ min normalised values and statistical significance evaluated using students t-test with Perseus. Volcano plots were generated using results from fold change analysis and t-test to display the differentially expressed metabolites based on statistical significance. Metabolite list available in table S1. Further pathway analysis was conducted using a hypergeometric test enrichment method and a relative-betweenness centrality topology measure according to the MetaboAnalyst 6.0 (KEGG) pathway library.

### Bacterial mutagenesis

*E. coli* mutant strains were constructed using lambda red homologous recombination as previously described(*64*) using the LR primers and plasmids listed in Supp Table 2. Correct unresolved insertions, antibiotic-resistance profiles, and subsequent resolved deletions after pCP20 transformation and flippase activity were verified using primers listed. For *in vivo* infections, fully resolved mutants were used to then make unresolved *lacZ* insertions as described above, but did not proceed to pCP20-mediated resolution. Strain list available in table S4.

### Mammalian cell line culture

CT26, HCT116, HCT116icasp9 & Caco2 were culture in DMEM + 10% FBS + L-glutamine (2mM) at 37°C, 5% CO_2_.

#### Apoptotic Supernatant Generation

HCT116/CT26 cells were seeded in 12 well plates (300,000 cells per well) in DMEM +10% FBS + L-Glut (2mM). Media was placed with DMEM (- additives) containing death induction trigger (staurosporine 1 μM or doxorubicin 20 μg mL^-1^, AP20187 2 pM) and the cells incubated for 6 or 24 h. Supernatant was removed, debris removed by centrifugation and filtered through a 0.22 μm filter. Apoptotic supernatants were aliquoted at 0.5 mL in 1.5 mL Eppendorf’s and stored at −80 °C until used. Viability of cells was ascertained via flow cytometry. For inhibitor treatment, cells were pretreated with the inhibitor for 60 min prior to addition of the death induction trigger. QVD (3 μM), dipyridamole (50 μM), NBMPR (1, 100, 250 μM).

#### Flow cytometry

For viability, cells were stained with annexin V (pacific blue or FITC, 1:500) and Draq7 (1:1000) for 30 min and immediately analysed by flow cytometry. Caspase 3/7 activity was analysed using the FLICA 660 caspase 3/7 activity kit (biorad; ICT9125). For antibody staining cells were pretreated with Trustain FCX (biorad, 1:1000). ENT-1/2 levels determined using ENT-1 Alexafluor 488 (abcam, 1:100) and ENT-2 Alexafluor 647 (abcam, 1:100). Analysis was carried out using Flojo v.10.

#### Apoptotic supernatant bacterial growth

All bacterial species/strains were cultured overnight in 3 mL of LB broth at 37 °C, 200 rpm. Apoptotic supernatants and controls were defrosted at 37 °C immediately prior to use. Bacteria were diluted to 1e^6^ CFU mL^-1^ and 1e^4^ bacteria added to the 0.5 mL aliquoted apoptotic supernatants. Bacteria were grown at 37 °C, 200 rpm for 7 h (staurosporine) or 16 h (doxorubicin) before dilution and plating on LB agar overnight. For anaerobic growth experiments, bacteria were sealed in an anaerobic jar containing anaerobic atmosphere generation bags (Merck). Colonies were counted and CFU mL^-1^ calculated.

#### Purine bacterial growth

Bacteria were cultured overnight in 3 mL of LB broth at 37 °C, 200rpm. DMEM was incubated at 37 °C immediately prior to use. Bacteria were diluted to 1e^6^ CFU mL^-1^ and 1e^4^ bacteria added to the 0.5 mL DMEM +/- selected purines. Bacteria were grown at 37 °C, 200rpm for 7h before dilution and plating on LB agar overnight. Colonies were counted and CFU mL^-1^ calculated.

#### Bacterial Complementation

The parent plasmid pGEN-MCS and pCA24N-ligase for bacterial complementation were a gift from Harry Mobley (Addgene plasmid #44919; RRID: Addgene #44919) and Wayne Patrick respectively (Addgene plasmid # 87741; RRID: Addgene_87741). Plasmid DNA purification was performed using the QIAprep spin miniprep kit (Qiagen), according to the manufacturer’s instructions. Primers were designed to amplify the open reading frames of the genes of interest (*purA*) from HS *E. coli* genomic DNA. For cloning into pGEN-MCS, the primers also amplified 500 bp upstream of *purA* to include their associated endogenous promoter (Supp Table 1). BamHI and NcoI restriction sites were also added to the 5’ and 3’ end of the oligonucleotides respectively. Platinum SuperFi II high-fidelity DNA polymerase (Thermo Fisher Scientific) was used for PCR amplification of the gene using HS *E. coli* genomic DNA as the template. The PCR product was purified using the QIAquick PCR purification kit (Qiagen). The respective vector and inserts were double digested with BamHI (New England Biolabs) and NcoI (New England Biolabs) restriction enzymes. The fragments were gel purified using QIAquick gel extraction kit (Qiagen) before overnight ligation with T4 DNA Ligase (New England Biolabs). The recombinant plasmids were transformed into DH5α electrocompetent *E. coli* (New England Biolabs) via electroporation, according to the suppliers’ instructions, and selected based on ampicillin (pGEN-MCS) or chloramphenicol (pCA24N-ligase) resistance. Successful constructs were verified by restriction enzyme digest, PCR, and Plasmidsaurus sequencing services. Verified clones were then transformed into the appropriate *HS E. coli* mutants via electroporation. Complementation was assessed by restoring the growth phenotype *in vitro.* Using complement pGEN-MCS, 1e^5^ of the bacteria were grown for 16 h with agitation in DMEM, supplemented with ampicillin, before diluting and plating on ampicillin resistant LB agar plates. For complement pCA24N-ligase, overnight growth was performed in LB broth supplemented with 25 mg ml^-1^ chlorophanicol +/- 0.4 mM IPTG. OD600s were read in a 1:10 dilution. Plasmid list available in Table S4.

#### Caspase 9 Activity

HCT116/CT26 cells were seeded in 12 well plates (300,000 cells per well) in DMEM +10% FBS + L-Glut (2mM). Media was placed with DMEM (- additives) containing death induction trigger (staurosporine 1 μM) and the cells incubated for 6 or 24 hours. Caspase 9 activity was measured following treatment using a Caspase Glo-9 Assay kit (Promega #G8210).

#### HCT116 inducible caspase 9 generation

The inducible caspase-9 (iCasp9) construct was adapted from the previously reported CharOFF-iCasp9 lentiviral expression vector (*65*). Specifically, the iCasp9 lentiviral vector was generated by amplifying the original vector backbone excluding the CharOFF element and re-circularizing via In-Fusion cloning (Takara Bio). The sequence of the re-circularized vector (pLV-SFFV-FKBP12(F36V)-inducible human Casp9-hPGK-Puro) was confirmed by next-generation sequencing (Plasmidsaurus).

Lentivirus was generated by transfecting Lenti-X 293T cells (Takara Bio) with pLV-SFFV-FKBP12(F36V)-inducible human Casp9-hPGK-Puro and the packaging plasmids psPAX2 (Addgene #12260) and pMD2.G (Addgene #12259) at a 6:8:1 ratio by mass. Transfection complexes were formed using TransIT-Lenti Transfection Reagent (Mirus) at a ratio of 3 μl TransIT-Lenti per 1 μg total DNA in Opti-MEM (Gibco). After 48 h, the lentiviral supernatant was collected and purified by filtration through a 0.45 μm SFCA filter. Purified lentiviral supernatants were then used to transduce HCT-116 cells (ATCC) as follows: 8 x 105 HCT-116 cells were plated in 2 ml growth media (DMEM supplemented with 10% heat-inactivated FBS, 1% Penicillin-Streptomycin-Glutamine) in 6-well tissue culture-treated plates and allowed to adhere overnight. The following day, 1 ml of fresh purified lentiviral supernatant was added together with 10 μg/ml polybrene (Millipore) to the adhered cells, which were then centrifuged at 800 g at 37°C for 90 min. Cells were placed in a 37°C/5% CO2 incubator and allowed to recover for 48 h. Antibiotic selection with 10 μg ml^-1^ puromycin (Thermo Fisher) was then carried out for 10 days. Puromycin-resistant cells were maintained as a polyclonal pool.

#### Scratch Assay

Caco2 cells were seeded in a 24 well plate (400,000 cells per well, DMEM, 10% FBS, L-Glut), pre annotated with 6 horizontal lines on the bottom of the wells, and grown at 37 °C, 5% CO_2,_ until confluent. Cells were scratched with a p20 pipette-tip vertically down the centre of the well crossing all 6 pre-marked lines. Media was replaced with either apoptotic supernatant from HCT116 icasp9 or DMEM +/- hypoxanthine. Scratch width was imaged using a Nikon TSR2 microscope on 40x magnification and measured using Fiji ImageJ, immediately after media change and after 24h. Supernatant treated wells were normalised to QVD treated cell supernatants. Hypoxanthine treated wells were normalised to untreated controls.

#### ATP Measurements

Caco2 cells were seeded in a white 96 well plate (50,000 cells per well, DMEM, 10% FBS, L-Glut) and grown at 37 °C, 5% CO_2_ for 24 h. Media was removed and replaced with DMEM + hypoxanthine (0, 10, 100, 1000 μM) for 0-2 h. Media was removed and the cells permeabilised with DMEM + tritonX-100 (1%) for 15 min. ATP measurements were carried out using the RealTime-Glo Extracellular ATP Assay (Promega #GA5010). Luminescence measurements were carried out on an Agilent Synergy Plate Reader H1.

#### Statistical analysis

For biological experiment analysis, data were analysed using prism 10 software (Graphpad), using the statistical analysis identified in figure legends. Data were assessed for outliers via ROUT (Q=0.1%), followed by testing for normality using the Shapiro-Wilk test. The appropriate parametric or nonparametric test is identified in respective figure legends.

---

**Figure S1.**
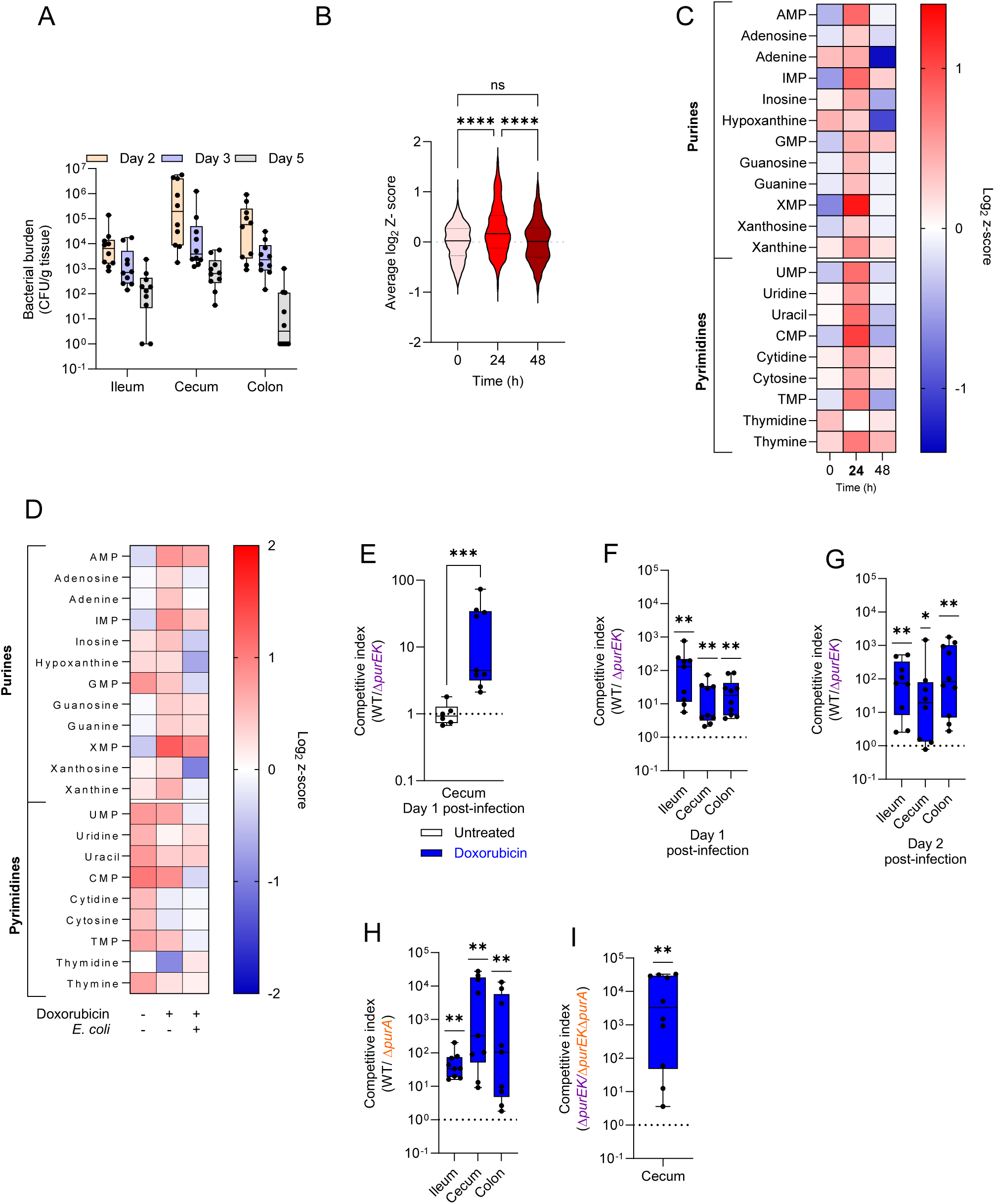
Purine utilisation by *E. coli in vivo*. **A**) *E. coli* HS growth kinetics in tissues up to 5 days post doxorubicin treatment and *E. coli* inoculation. **B**) Aggregated metabolite log_2_-Z-scores from faecal pellets at 0, 24 and 48h post doxorubicin treatment. n = 389 metabolites, one-way Anova. **C**) Relative purine and pyrimidine levels in faecal pellets at 0, 24 and 48h post doxorubicin treatment. **D**) Relative purine and pyrimidine levels in faecal pellets at 24h from untreated, doxorubicin treated and doxorubicin treated-*E. coli* inoculated mice. **E**) Competitive index of *E. coli* HS WT (SR007) vs *ΔpurEK* (PC055) in cecum 1 day post infection (dpi) in untreated (white) and doxorubicin treated (blue) mice. n ≥ 6, unpaired t-test **F**) Competitive index of *E. coli* HS WT (SR007) vs *ΔpurEK* (PC055) in ileum, cecum & colon 1 dpi & **G**) 2 dpi in doxorubicin treated mice. n > 9, one sample Wilcoxon test. **H**) Competitive index of *E. coli* HS WT (SR007) vs *ΔpurA* (PC118) in ileum, cecum & colon 2 dpi in doxorubicin treated mice. n = 10, one sample Wilcoxon test. **I**) Competitive index of *E. coli* HS *ΔpurEK* (PC055) vs *ΔpurEK-purA* (PC120) in cecum 2 dpi in doxorubicin treated mice. n = 10, one sample Wilcoxon test.

**Figure S2.**
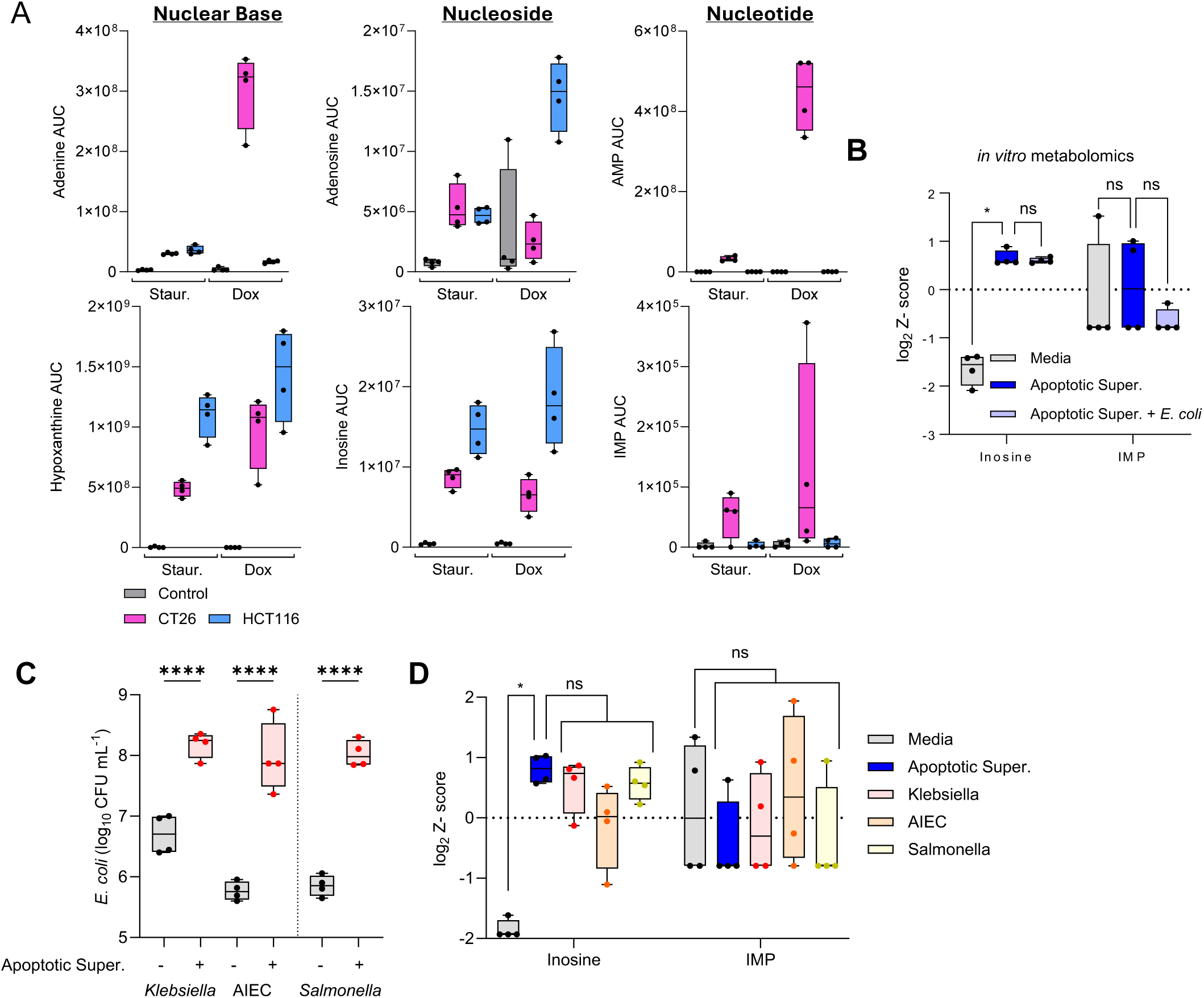
**Hypoxanthine is preferentially utilised by Enterobacteriaceae**. **A**) Levels of purines found in apoptotic supernatants from CT26 and HCT116 cells treated with staurosporine (1 μM) or doxorubicin (20 μg mL^-1^) for 24h. n = 4. **B**) Log_2_ Z-score of inosine and IMP extracted from fig 2H. n = 4, one-way Anova. **C**) CFU mL^-1^ after 7h of *K. pneumoniae*, *E. coli* LF82 (AIEc) & *S. enterica* SL1344 grown in media controls or apoptotic supernatants (HCT116 cells, staurosporine (1 μM), 6h) . n = 4, one-way anova. **D**) Log_2_ Z-score of inosine and IMP in apoptotic supernatants inoculated with *K. pneumoniae*, *E. coli* LF82 & *S. typhimarium* SL1344. n=4, one-way Anova.

**Figure S3.**
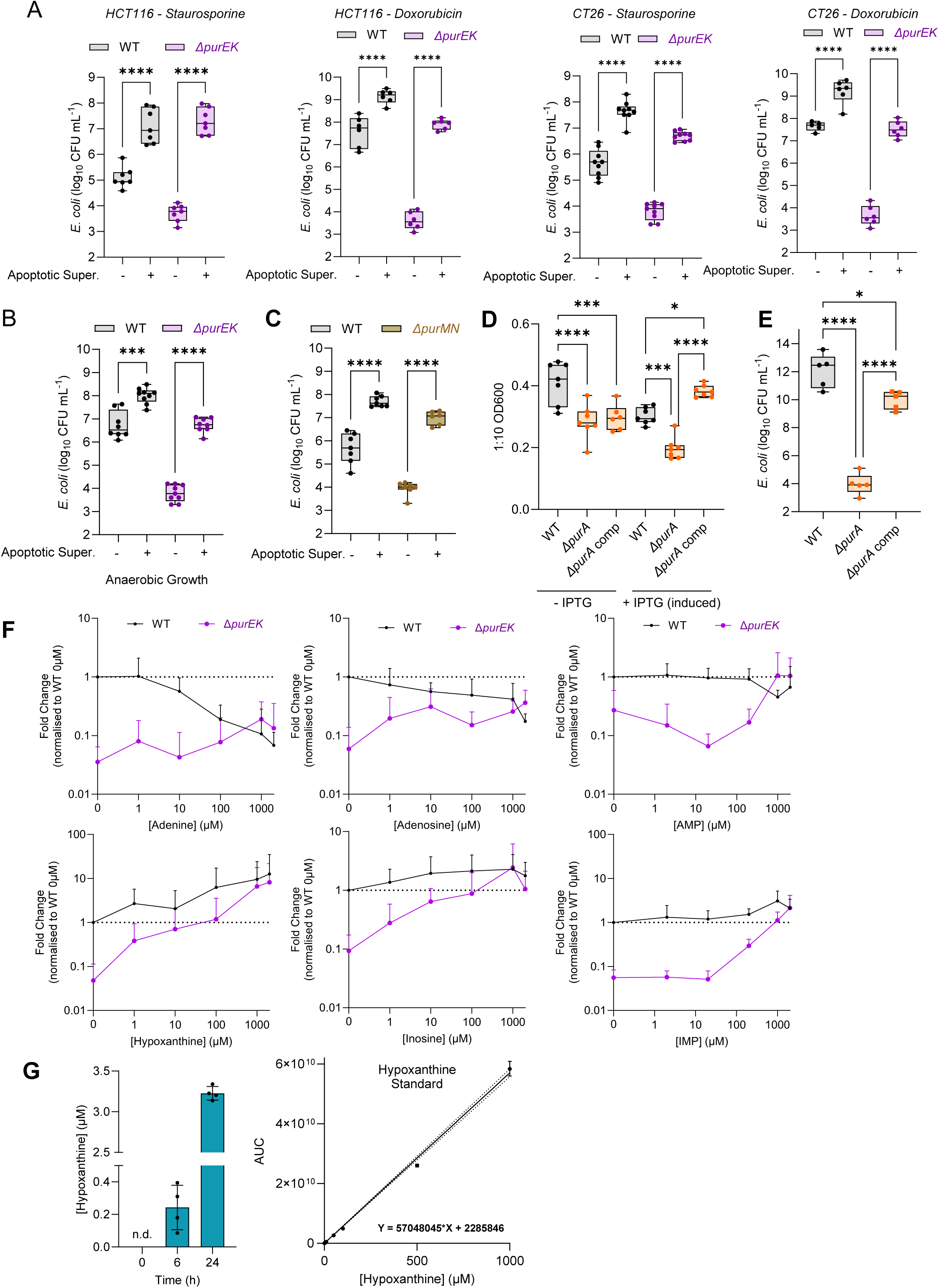
***E. coli* salvage purines *in vitro*. A**) CFU mL^-1^ of *E. coli* HS WT (grey) vs *ΔpurEK* (CJA117, purple) grown for 7h in media controls or apoptotic supernatants from CT26 or HCT116 cells treated for 6h with doxorubicin (20 μg mL^-1^) or staurosporine (1 μM). n ≥ 5, one-way Anova. **B**) CFU mL^-1^ of *E. coli* HS WT (grey) vs *ΔpurEK* (purple) grown under anaerobic conditions for 7h in media controls or apoptotic supernatants (HCT116 cells, Sstaurosporine (1 μM), 6h) . n ≥ 7, one-way Anova. **C**) CFU mL^-1^ of *E. coli* HS WT (grey) vs *ΔpurMN* (CJA128, brown) grown for 7h in media controls or apoptotic supernatants (CT26 cells, staurosporine (1 μM), 6h) n >6, one-way Anova. **D**) OD_600_ of *E. coli* HS WT (PC179), *ΔpurA* (PC184) *& ΔpurA* complemented with *purA*-pCA24N (PC192) +/- IPTG induction (0.4 mM) for 16h. n>6, one-way Anova. **E**) Growth of *E. coli* HS WT (PC234, grey) vs *ΔpurA* (PC227, orange circles) vs *ΔpurA* complemented with *purA*-pGEN-MCS purA plasmid (PC187, orange squares). n = 5, one-way Anova. **F**) Growth of *E. coli* HS WT (grey) vs *ΔpurEK* (CJA117, purple) grown in media containing stated purines for 7h. **G**) Quantification of hypoxanthine in supernatants taken from HCT116 cells treated with staurosporine. n=4. Standard curve for the determination of hypoxanthine concentrations in apoptotic supernatants. n = 3.

**Figure S4.**
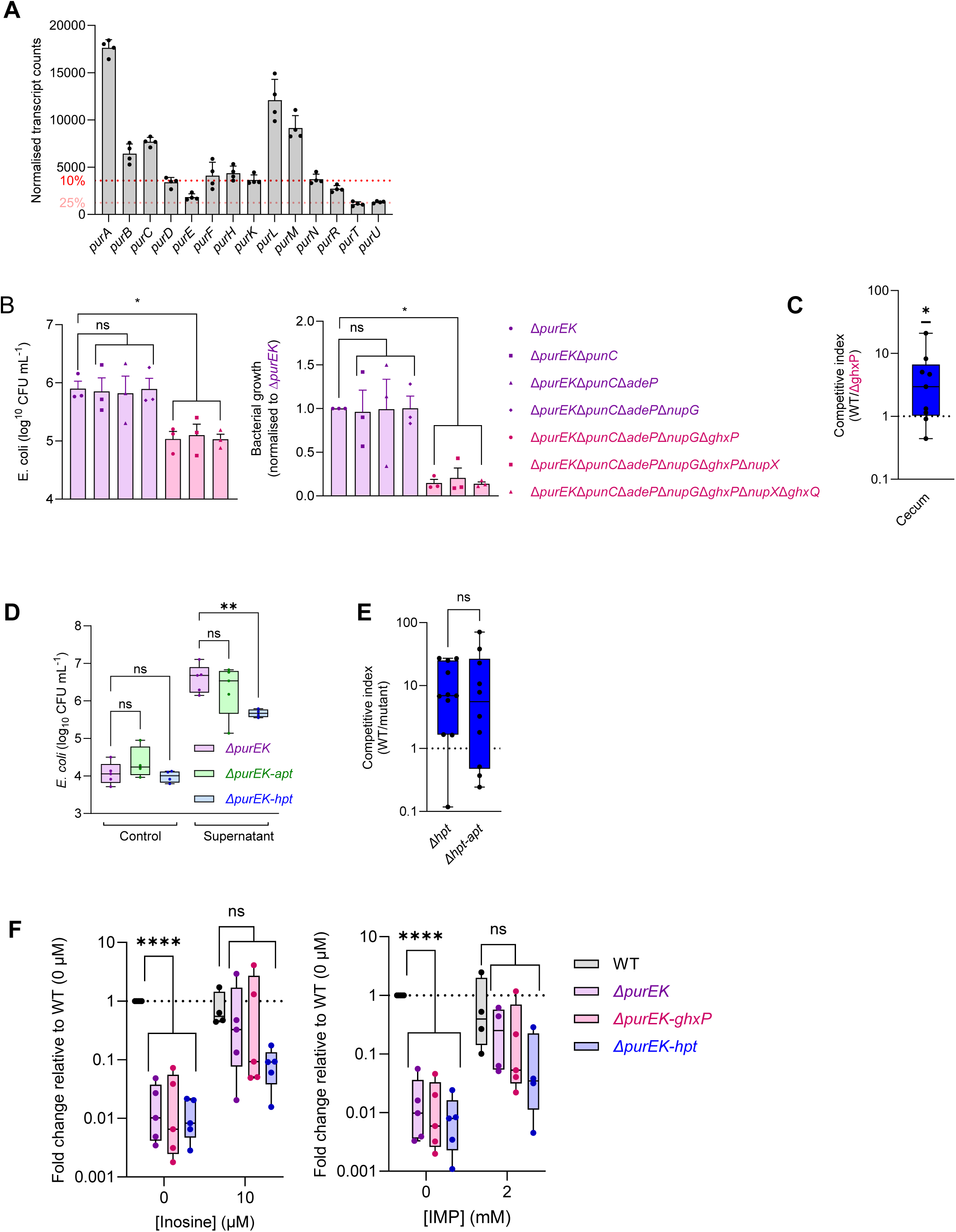
Hypoxanthine transport and salvage are gene specific. **A**) Transcript counts for genes in the purine biosynthesis and salvage chains of *E. coli* HS. Lines indicate values in the top 10% (red) and 25% (pink) of counts. n = 4. **B**) CFU mL^-1^ of *E. coli* HS *ΔpurEK* vs *ΔpurEK* + alternative purine transporters (CJA117, PC056, SR040, SR052, PC125, PC136, PC146, PC054) grown for 7h in apoptotic supernatants (HCT116 cells, Staurosporine (1 μM), 6h). n = 3, one-way Anova. **C**) Competitive index of *E. coli* HS WT (SR007) vs *ΔghxP* (PC230) in cecum 2 dpi in doxorubicin treated mice. n = 9, one sample Wilcoxon test. **D**) CFU mL^-1^ of *E. coli* HS *ΔpurEK* (CJA117, purple) vs *ΔpurEK-apt* (SJB040, green) & *ΔpurEK-hpt* (SJB044, blue) grown for 7h in media controls or apoptotic supernatants. (HCT116 cells, Staurosporine (1 μM), 6h). n ≥ 4, one-way Anova. **E**) Competitive index between *E. coli* HS WT (SR007) and *Δhpt* (PC169) or *Δhpt-apt* (PC175) in faecal pellets 2 dpi. n ≥ 9, Mann-Whitney test. **F**) CFU mL^-1^ of *E. coli* HS WT, *ΔpurEK* (CJA117, purple)*, ΔpurEK-ghxP* (PC177, pink) & *ΔpurEK-hpt* (SJB044) grown in media +/- IMP or inosine. n >4, one-way Anova.

**Figure S5.**
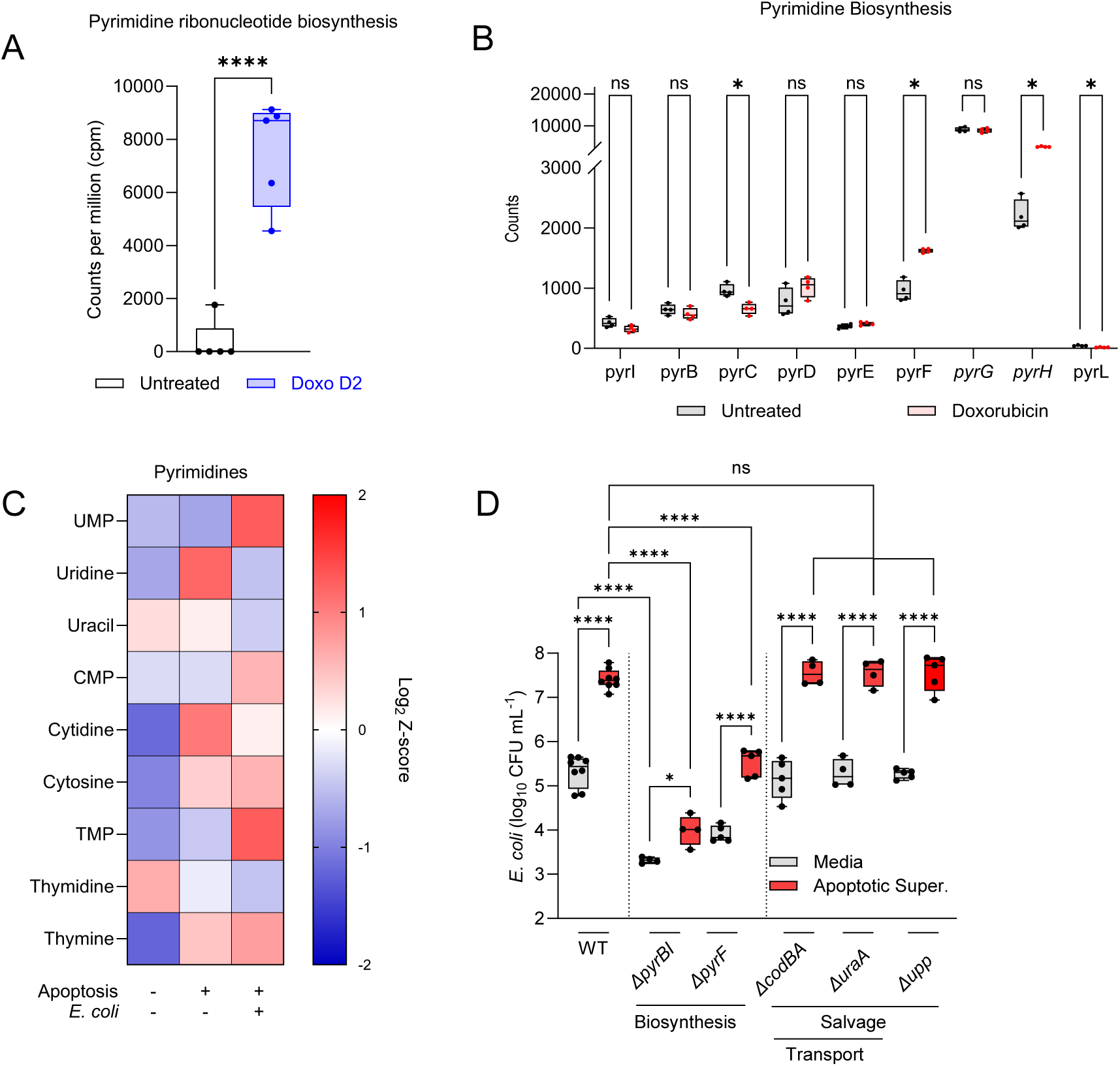
**Pyrimidine salvage is not sufficient to maintain growth**. **A**) Meta transcriptomics counts from intestinal microbiota taken from doxorubicin treated mice (d2). n = 5, unpaired t-test. **B**) *E. coli* transcript counts for pyrimidine biosynthesis, transport & salvage genes. n = 4, multiple t-test. **C**) Relative pyrimidine levels in media controls, 6h staurosporine (1 μM) treated HCT116 apoptotic supernatants and 6h staurosporine (1 μM) treated HCT116 apoptotic supernatants inoculated with *E. coli* HS for 7h. n = 4. **D**) CFU mL^-1^ of *E. coli* HS WT vs pyrimidine biosynthesis and salvage mutants (PC243, SJB052, SR011, PC251, PC249) grown for 7h in media or apoptotic supernatants. (HCT116 cells, Staurosporine (1 μM), 6h). n ≥ 4, one-way Anova.

**Figure S6.**
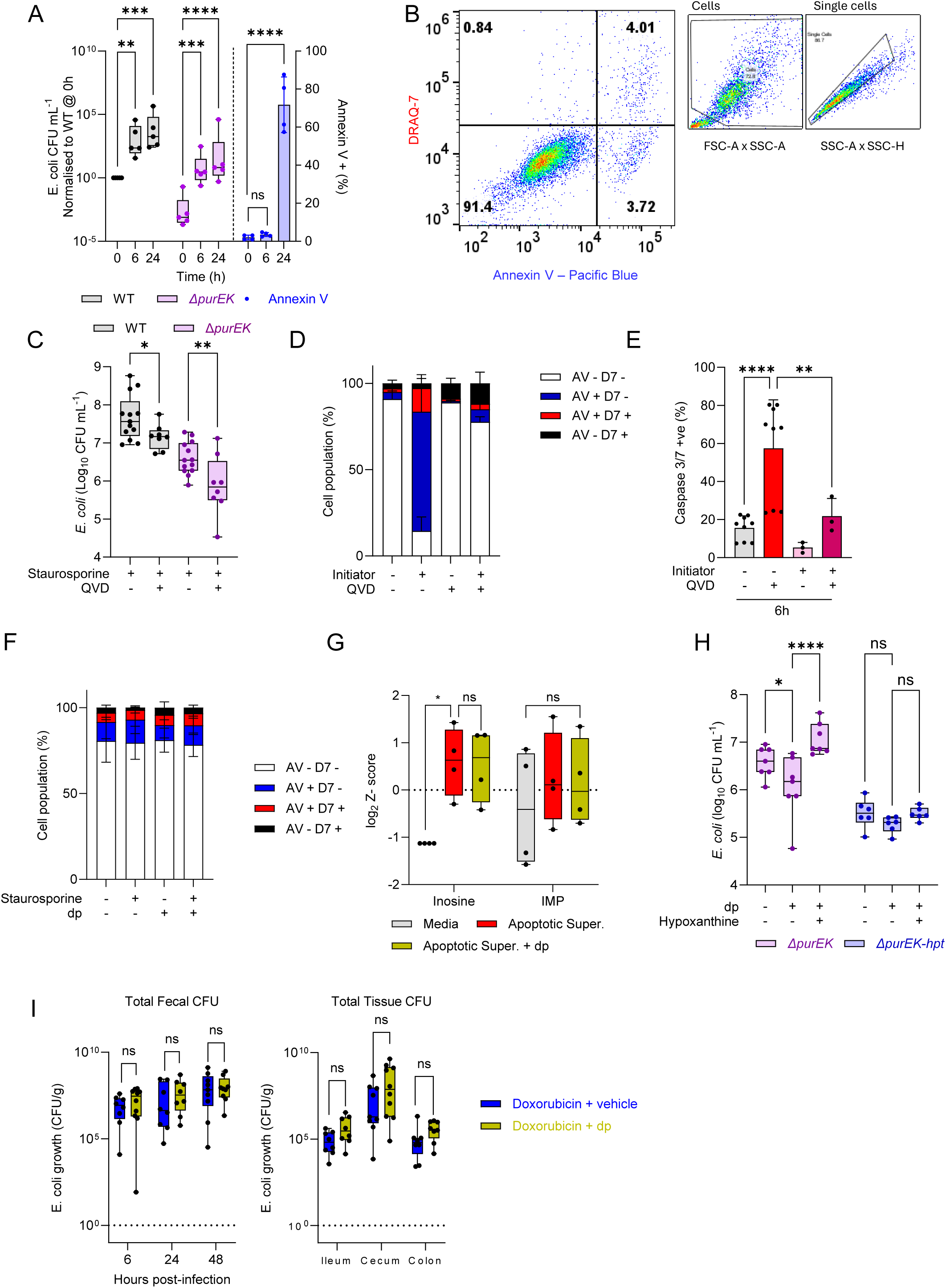
Purine release originates in early apoptosis. **A**) CFU mL-1 of E. coli HS WT (grey) vs *ΔpurEK* (CJA117, purple) grown for 7h in media controls or apoptotic supernatants (CT26 cells, staurosporine (1 μM), 6/24h) n ≥ 4, one-way Anova. Alongside the levels of Annexin V (blue) staining in HCT116 cells treated for 0, 6 or 24h with staurosporine. n = 9, one-way Anova. **B**) Gating strategy to determine Annexin V and Draq7 positivity. Cell viability of HCT116 *icasp9* cells treated with initiator AP20187 (2 pM) for 6h +/- pretreatment with QVD (3 uM) via **C**) CFU mL^-1^ of E. coli HS WT (grey) vs *ΔpurEK* (CJA117, purple) grown for 7h in media controls or apoptotic supernatants (HCT116 cells, staurosporine (1 μM), 6h, +/- QVD (3 μM)). n ≥ 8, one-way Anova. **D**) Annexin V (pacific blue) and draq7 staining (n = 4) or **E**) caspase 3/7 activity (n ≥ 3, one-way Anova) in HCT116 cells treated with initiaitor +/- QVD for 6h. **F**) Annexin V (FITC) and draq7 staining of HCT116 cells treated +/- staurosporine (1 μM) and +/- dp (50 μM). n = 4. **G**) Relative levels of purines in supernatants taken from HCT116 cells treated +/- staurosporine (1 μM) and +/- dp (50 μM). n = 4, one-way Anova. **H**) CFU mL-1 of E. coli HS *ΔpurEK* (CJA117, purple) vs *ΔpurEK-hpt* (SJB044, blue) grown for 7h in media controls or apoptotic supernatants (HCT116 cells, staurosporine (1 μM), 6h) +/- pretreatment with dp (50 μM) +/- subsequent addition of hypoxanthine (10 μM). n ≥ 4, one-way Anova. **I**) Total faecal CFU of mice treated with doxorubicin (15 mg kg^-1^) +/- dp (30 mg kg^-1^) at 6, 24 and 48h post doxorubicin treatment & total tissue CFU at 48h post doxorubicin treatment. n ≥ 7, multiple Mann-Whitney tests.

**Figure S7.**
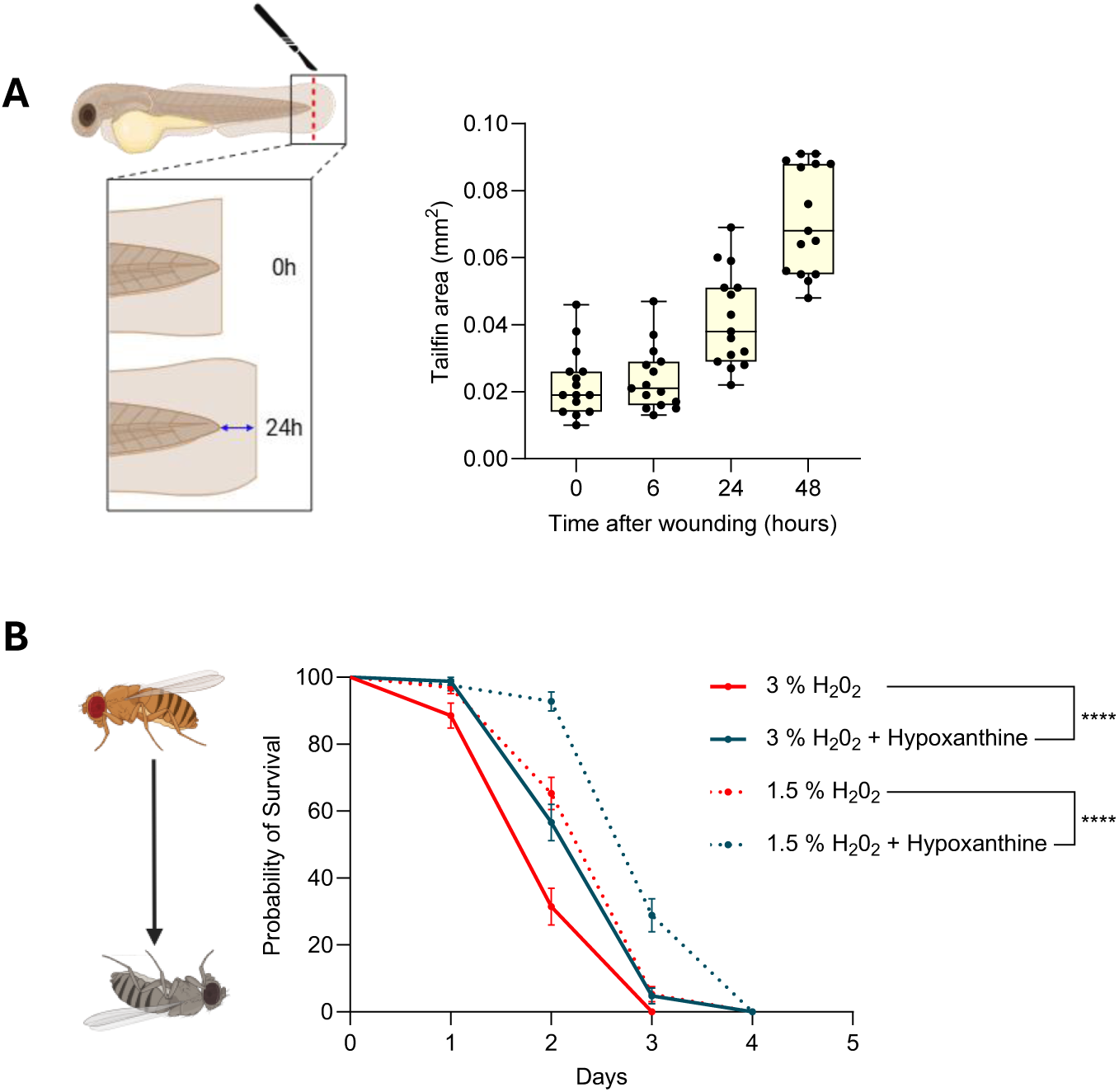
**A**) Regrowth of resected tail fin over 48h in untreated zebrafish embryos. n > 15. **B**) Survival curve of *Drosophila* treated +/- H_2_O_2_, +/- hypoxanthine (100 μM). n >70, Kaplan-Meier test.

