## Supplemental Tables for "Intestinal bacteria hijack an evolutionarily conserved epithelial repair signal"

| Table S1 – Metabolites analysed by metabolomics. | | |
|---|---|---|
| (2E,4E)-2,4-Hexadienoic acid | 1-Aminocyclopropanecarboxylic acid | 1-Methyladenosine |
| 1-Methylhistidine | 2,3-Diaminopropionic acid | 2',4'-Dihydroxyacetophenone |
| 2,4-dihydroxypteridine | 2,6-Dihydroxypyridine | 2-Amino-2-deoxy-D-gluconate |
| 2-Aminoisobutyric acid | 2-Aminophenol | 2-hydroxy-4-(methylthio)butanoate |
| 2-Hydroxybutyric acid | 2-Hydroxyphenethylamine | 2-hydroxyphenylacetate / mandelate |
| 2-Hydroxypyridine | 2-Keto-3-deoxy-D-gluconic acid | 2-Methylcitric acid |
| 2-Methylglutaric acid | 2-Phosphoglyceric acid | 2-Pyrocatechuic acid |
| 3-(2-Hydroxyphenyl)propanoic acid | 3,4-Dihydroxybenzeneacetic acid | 3,4-Dihydroxymandelic acid |
| 3,4-Dihydroxyphenylglycol | 3-Amino-4-hydroxybenzoate | 3-Dehydroshikimate |
| 3-Hydroxyanthranilic acid | 3-Hydroxybenzoic acid | 3-Hydroxybenzyl alcohol |
| 3-Hydroxybutyric acid | 3-Hydroxymethylglutaric acid | 3-Hydroxyphenylacetic acid |
| 3-Methoxytyramine | 3-Methoxytyrosine | 3-methyl-2-oxindole |
| 3-Methyl-2-oxovaleric acid | 3-Methyladenine | 3-Methylglutaconic acid |
| 3-Methylhistamine | 3-Methylhistidine | 3-Phosphoglyceric acid |
| 3-Sulfinoalanine | 4,5-Dihydroorotic acid | 4-Guanidinobutanoic acid |
| 4-Hydroxybenzaldehyde | 4-Hydroxybenzoic acid | 4-Hydroxycinnamic acid |
| 4-Hydroxynonenal | 4-Hydroxyproline | 4-Methylcatechol |
| 5,6 dimethylbenzimidazole | 5-Aminolevulinic acid | 5-Aminopentanoic acid |
| 5a-Tetrahydrocortisol | 5'-Deoxyadenosine | 5-Hydroxyindoleacetic acid |
| 5-Hydroxy-L-tryptophan | 5-Methylcytosine | 5'-Methylthioadenosine |
| 5-Thymidylic acid | 6-Hydroxynicotinic acid | 6-Methylthiopurine |
| 6-Phosphogluconic acid | Acetoacetic acid | Acetylcholine chloride |
| Acetyl-CoA | Acetylcysteine | Acetylglycine |
| Adenine | Adenosine | Adenosine 2',3'-cyclic phosphate |
| Adenosine 3',5'-diphosphate | Adenosine monophosphate | Adenosine triphosphate |
| Adipic acid | ADP | ADP-glucose |
| ADP-ribose | Agamatine sulfate | AICAR |
| Allantoin | Allose | Alpha-D-Glucose |
| Aminoadipic acid | aminoisobutanoate | aniline-2-sulfonate |
| Anserine | arabinose/ribose/xylose/lyxose | arabitol/ribitol/xylitol |
| Ascorbate | Asymmetric dimethylarginine | Azelaic acid |
| Benzaldehyde | Benzyl alcohol | Benzylamine |
| beta-Alanine | Beta-Glycerophosphoric acid | Betaine |
| Biotin | butiric acid | Caffeate |
| Carbomyl Phosphate | Carnosine | CDP |
| CDP-Ethanolamine | Choline | Ciliatine |
| cis-4-Hydroxy-D-proline | Citicoline | Citramalic acid |
| Citric acid | Citrulline | Coenzyme A |
| Cortisone | Creatine | Creatinine |
| Cyclic AMP | Cyclic GMP | Cysteamine |
| Cysteic acid | Cysteinylglycine | Cytidine |
| Cytidine 2',3'-cyclic phosphate | Cytidine monophosphate | Cytidine Triphosphate |
| Cytosine | D-Alanine | dCDP |
| dCMP | Deoxyadenosine | Deoxyadenosine monophosphate |
| Deoxyadenosine triphosphate | Deoxycytidine | Deoxyguanosine |
| Deoxyuridine | Deoxyuridine triphosphate | Dethiobiotin |
| dGDP | dGMP | dGTP |
| Diacetyl | Diaminopimelic acid | Diethanolamine |
| Dihydrobiopterin | Dihydrofolic acid | Dihydrouracil |
| Dihydroxyacetone phosphate | Dimethylallylpyrophosphate | DL-Homocystine |
| D-Ribose 5-phosphate | dTDP-glucose | dUMP |
| Epinephrine | Erythritol | Ethylmalonic acid |
| FAD | Folic acid | Fructose 1,6-bisphosphate |
| Fructose 6p/ galactose 1p/mannose 6p | Fumaric acid | Galactaric acid |
| Galactitol | Galactosamine | Galactose 1-phosphate |
| galactose/tagatose/mannose/sorbose/psicose/myoinositol | Galacturonic acid | Geranyl-PP |
| Glucaric acid | Gluconic acid | Gluconolactone |
| Glucosamine | Glucosamine 6-phosphate | Glucosamine 6-sulfate |
| Glucose 1-phosphate | Glucose 6-phosphate | Glucuronic acid |
| Glucuronolactone | Glutaric acid | Glutathione |
| Glyceraldehyde | Glyceric acid | Glycerol |
| Glycerol 3-phosphate | Glycine | Glycolate |
| Guanidinosuccinic acid | Guanidoacetic acid | Guanine |
| Guanosine | Guanosine diphosphate | Guanosine diphosphate mannose |
| Guanosine monophosphate | Guanosine triphosphate | Hippuric acid |
| Histamine | Homocysteine | Homogentisic acid |
| Homovanillic acid | Hydroquinone | Hydroxykynurenine |
| Hydroxyphenyllactic acid | Hypotaurine | Hypoxanthine |
| IDP | Imidazoleacetic acid | Indoleacetaldehyde |
| Indoleacetic acid | indoxyl b-glucoside | Indoxyl sulfate |
| Inosine | Inosinic acid | Isocitric acid |
| Isopentenyl pyrophosphate | Itaconic acid | ITP |
| Ketoleucine | Kynurenic acid | lactose/melibiose/sucrose/cellobiose/maltose/palatinose |
| L-Alanine | L-Allothreonine | L-Arginine |
| L-Asparagine | L-Aspartic acid | L-Cystathionine |
| L-Cysteine | L-Cystine | L-Dehydroascorbic acid |
| L-Dopa | L-Glutamic acid | L-Glutamine |

| | | |
|---|---|---|
| L-Gulonolactone | L-Gulose | L-Histidine |
| L-Histidinol | L-Homocysteine thiolactone | L-Homoserine |
| Lipoamide | L-Isoleucine | L-Kynurenine |
| L-Lactic acid | L-Leucine | L-Lysine |
| L-Malic acid | L-Methionine | L-Norleucine |
| L-Phenylalanine | L-Proline | L-Threonine |
| L-Tryptophan | L-Tyrosine | L-Valine |
| Maleic acid | Malonate | Malondialdehyde |
| mannitol/sorbitol | meso-Tartaric acid | methyl 4-aminobutyrate |
| Methyl beta-D-glucopyranoside | Methylmalonic acid | Mevalonic acid |
| Monoethyl malonic acid | N6,N6,N6-Trimethyl-L-lysine | N-Acetylasparagine |
| N-Acetyl-D-glucosamine | N-Acetylgalactosamine | N-Acetylglutamic acid |
| N-Acetyl-L-aspartic acid | N-Acetylleucine | N-Acetylmannosamine |
| N-Acetylmethionine | N-Acetylneuraminate | N-Acetylproline |
| N-Acetylputrescine | N-Acetylserine | N-Acetylserotonin |
| NAD | NADH | NADP |
| NADPH | N-Alpha-acetyllysine | N-Formylglycine |
| N-Formyl-L-methionine | Niacinamide | Nicotinamide hypoxanthine dinucleotide |
| Nicotinamide ribotide | Nicotinic acid | N-Methylalanine |
| N-methyl-L-glutamic Acid | N-Methyltryptamine | Norepinephrine |
| Normetanephrine | o-Methoxyphenol | O-Phosphoethanolamine |
| Ophthalmic acid | Ornithine | Orotic acid |
| O-Succinyl-L-homoserine | Oxalacetic acid | Oxalic acid |
| Oxidized glutathione | Oxoadipic acid | Oxoglutaric acid |
| p-Aminobenzoic acid | Pantothenic acid | Paraxanthine |
| Phenol | Phenylpyruvic acid | Phosphocreatine |
| Phosphoenolpyruvic acid | Phosphonoacetate | Phosphorylcholine |
| Phosphoserine | p-Hydroxyphenylacetic acid | Picolinic acid |
| Pimelic acid | Pipecolic acid | Pivalic acid |
| p-Octopamine | Porphobilinogen | procollagen 5-hydroxy-L-lysine |
| Protocatechuic acid | Pterin | Purine |
| Pyridoxal | Pyridoxamine | Pyridoxine |
| Pyrocatechol | Pyroglutamic acid | Pyrrole-2-carboxylic acid |
| Pyruvic acid | Quinaldic acid | Quinic acid |
| Quinolinic acid | Raffinose | Retinol |
| Rhamnose | S-Adenosylhomocysteine | S-Adenosylmethionine |
| Salicylamide | Salsolinol | Sarcosine |
| Sebacic acid | Sedoheptulose | Serine |
| Serotonin | Shikimic acid | Sphinganine |
| Stachyose | Suberic acid | Succinic acid |
| Tartaric acid | Taurine | Tetrahydrofolic acid |
| Theobromine | Thiamine | Threitol |
| Thymidine | Thymine | Thyrotropin releasing hormone |
| Thyroxine | trans-Aconitic acid | trans-Cinnamate |
| trans-Ferulic acid | Trehalose | Trigonelline |
| Tryptamine | Tryptophanamide | Tyramine |
| Uracil | uracil 5-carboxylate | Ureidopropionic acid |
| Uric acid | Uridine | Uridine 5'-diphosphate |
| Uridine 5'-monophosphate | Uridine diphosphate glucuronic acid | Uridine diphosphategalactose |
| Uridine diphosphate-N-acetylgalactosamine | Uridine diphosphate-N-acetylglucosamine | Uridine triphosphate |
| Urocanic acid | Vanillylmandelic acid | Vanylglycol |
| Xanthine | Xanthosine | Xanthurenic acid |
| Xanthylic acid | | |

| Table S2. Primers | | | | | |
|---|---|---|---|---|---|
| *Gene* | *EcHS* | *Primer* | *Mutagenesis /Sequencing /Cloning* | *Forward /Reverse* | *Primer Sequence* |
| purEK | A0598/ A0597 | purE_LR_F1 | M | F | AGACGCATGTCTTCCCGCAATAATCCGGCGCGTGTCGCCATCGTGATGGGGTGTAGGCTGGAGCTGCTTC |
| | | purK_LR_R1 | M | R | CAGGTTAACTGAACTTACTCTGTGCCCACATCACGCCGCTGGCATATTCCCATATGAATATCCTCCTTAG |
| | | purE_F1 | S | F | CCTGGCATACGGAAGGTTCA |
| | | purK_R1 | S | R | AAGCGGGGACCTGTATTTCG |
| purA | A4420 | purA_LR_F1 | M | F | TGAAAAATGGGTAACAACGTCGTCGTACTGGGCACCCAATGGGGTGACGAGTGTAGGCTGGAGCTGCTTC |
| | | purA_LR_R1 | M | R | AGAATTACGCGTCGAACGGATCGCGCAGAATCATGGTTTCAGTACGATCCCATATGAATATCCTCCTTAG |
| | | purA_F1 | S | F | TTTACGTCGTTTTGGCGGTG |
| | | purA_R1 | S | R | ATGGCTTAACCGACACAGCA |
| purMN | A2634 / A2635 | purM_LR_F1 | M | F | CAAGCAGTGACCGATAAAACCTCTCTTAGCTACAAAGATGCCGGTGTTGAGTGTAGGCTGGAGCTGCTTC |
| | | purN_LR_R1 | M | R | GGCATTACTCGTCGGCAGCGTAGCCCTGCGGCGGCAGACGTTGACCATCCCATATGAATATCCTCCTTAG |
| | | purM_F1 | S | F | TACAAGTCTTCTTTCGCCGC |
| | | purN_R1 | S | R | CGGGACACTGCCTGGATATT |
| apt | A0546 | apt_LR_F1 | M | F | ACACTTATGACCGCGACTGCACAGCAGCTTGAGTATCTCAAAAATAGCATGTGTAGGCTGGAGCTGCTTC |
| | | apt_LR_R1 | M | R | ATAATTAATGGCCCGGGAACGGGACAAGGCTGTAGCTGGTAATGCCCTGTCATATGAATATCCTCCTTAG |
| | | apt_F1 | S | F | TGGCGAATTCCGGTGATTGA |
| | | apt_R1 | S | R | CGTCAGCAAAGGTTTGTGGG |
| hpt | A0129 | hpt_LR_F1 | M | F | AGAGATATGAAACATACTGTAGAAGTAATGATCCCCGAAGCGGAGATTAAGTGTAGGCTGGAGCTGCTTC |
| | | hpt_LR_R1 | M | R | TCACACTTACTCGTCCAGCAGAATCACTTTGCCGATATACGGCAGATGACCATATGAATATCCTCCTTAG |
| | | hpt_F1 | S | F | GTAATCGTCGCGAGCCTGTA |
| | | hpt_R1 | S | R | AGTTACCATTCACGGCTGGG |
| ghxP | A4307 | ghxP_LR_F1 | M | F | CCACGCATCTTGACGAAAATTAAACTCTCAGGGGATGTTTTCTTATGTCTGTGTAGGCTGGAGCTGCTTC |
| | | ghxP_LR_R1 | M | R | AAGATTAGATAGCCCAGCCACCCGCATAGAAGGTCACCAGCGCCACGGCGCATATGAATATCCTCCTTAG |
| | | ghxP_F1 | S | F | CCGCTATGATCGCCGGATAA |
| | | ghxP_R1 | S | R | TTCCATGCGTTCCCTGTTTG |
| ghxQ | A3044 | ghxQ_LR_F1 | M | F | CGCAACTAATAATAAAACTTTAACATCCTCGTGAGGACATCATTATGTCTGTGTAGGCTGGAGCTGCTTC |

| | | | | | |
|---|---|---|---|---|---|
| | | ghxQ_LR_R1 | M | R | TCTATTAGATTGCCCAACCACCCGCGTAAAATGCGACCAGTGCGGCAGTACATATGAATATCCTCCTTAG |
| | | ghxQ_F1 | S | F | CTACAGCAGCTGCGCTATGA |
| | | ghxQ_R1 | S | R | CCAGCCTGCGTTGAATCTTG |
| punC | A1740 | punC_LR_F1 | M | F | CACTTTGCGGCGGTGTTTAATTGAGAGATTTAGAGAATATACATGCAACCGTGTAGGCTGGAGCTGCTTC |
| | | punC_LR_R1 | M | R | CAACAAAATTTTGTTAGCAGCCTAAGTATAAGTATATCGATATAGATCAGCATATGAATATCCTCCTTAG |
| | | punC_F1 | S | F | TGACGCACGGTATAGCTGAC |
| | | punC_R1 | S | R | AACCCTTGCCGAAATGACCA |
| adeP | A3928 | adeP_LR_F1 | M | F | TATACGATAGCGACGGATTTTCCCTCCTTGTTCGGAAATAGATAATGAGTGTGTAGGCTGGAGCTGCTTC |
| | | adeP_LR_R1 | M | R | GATTTTAATGAGCGTCGATAAATACAATCTTCAGGATAAACAGCAGCGCACATATGAATATCCTCCTTAG |
| | | adeP_F1 | S | F | CGCAGTCGAAAAATACCGCT |
| | | adeP_R1 | S | R | TTGATCCGCAAACCGGAGAA |
| nupG | A3125 | nupG_LR_F1 | M | F | ATTAACATGAATCTTAAGCTGCAGCTGAAAATCCTCTCTTTTCTGCAGTTGTGTAGGCTGGAGCTGCTTC |
| | | nupG_LR_R1 | M | R | GTAATTAGTGGCTAACCGTCTGTGTGCCTGTCGGGACACGAACGTGTTTACATATGAATATCCTCCTTAG |
| | | nupG_F1 | S | F | ATTAACCGCCCCTGACGATG |
| | | nupG_R1 | S | R | GCACTGGAAGAAGGGGTGAA |
| nupX | A2297 | nupX_LR_F1 | M | F | ATAACTATGGATGTCATGAGAAGTGTTCTGGGAATGGTGGTATTGCTGACGTGTAGGCTGGAGCTGCTTC |
| | | nupX_LR_R1 | M | R | CGCACTACGCTAAACCAATAAAGAATCCTGCAATAGTAGCACTCATCAGGCATATGAATATCCTCCTTAG |
| | | nupX_F1 | S | F | GGCTTAGTCGAAGAGTGCGT |
| | | nupX_R1 | S | R | AGGATACAGTTTCCGCGACA |
| pyrBI | A4500 / A4501 | pyrB_LR_F1 | M | F | TAAAAGATGGCTAATCCGCTATATCAGAAACATATCATTTCCATAAACGAGTGTAGGCTGGAGCTGCTTC |
| | | pyrI_LR_R1 | M | R | GCAATTAATTGGCCAGCACCACATTATGGGAAAACTCTTTTTCACAGTATCATATGAATATCCTCCTTAG |
| | | pyrB_F1 | S | F | TTGATCACCCATTCCCAGCC |
| | | pyrI_R1 | S | R | GCCCGTTTTCGGATTTACCG |
| pyrF | A1393 | pyrF_LR_F1 | M | F | CTGGTCATGACGTTAACTGCTTCATCTTCTTCCCGCGCTGTTACGAATTCGTGTAGGCTGGAGCTGCTTC |
| | | pyrF_LR_R1 | M | R | CTCATCATGCACTCCGCTGTAAAGAGGCGTTGATCGCTTTCAGCGTCTGCCATATGAATATCCTCCTTAG |
| | | pyrF_F1 | S | F | ATGCTCGCCGTTTACCTGTT |
| | | pyrF_R1 | S | R | AGGCAAACGCCCTTACCTTT |
| upp | A2633 | upp_LR_F1 | M | F | AAGAGTATGAAGATCGTGGAAGTCAAACACCCACTCGTCAAACACAAGCTGTGTAGGCTGGAGCTGCTTC |
| | | upp_LR_R1 | M | R | TTCTTTATTTCGTACCAAAGATTTTGTCACCGGCATCGCCGAGGCCCGGACATATGAATATCCTCCTTAG |

| | | | | | |
|---|---|---|---|---|---|
| | | upp_F1 | S | F | CGAAAGAAGACTTGTACCAGGG |
| | | upp_R1 | S | R | GCACCAAACATGGCGAACAA |
| uraA | A2632 | uraA_LR_F1 | M | F | AATACTATGACGCGCCGTGCTATCGGGGTGAGTGAAAGACCGCCACTTTTGTGTAGGCTGGAGCTGCTTC |
| | | uraA_LR_R1 | M | R | TACATTATTTGTCTGTTATATCCGCGTCTTCTGCGTCCAGCACCACTTCTCATATGAATATCCTCCTTAG |
| | | uraA_F1 | S | F | CCGCATCGATTGATCAGGGA |
| | | uraA_R1 | S | R | GCGGCCAGTAAAGAGGAGTT |
| codBA | A0401 | codB_LR_F1 | M | F | AATTTCGTGTCGCAAGATAACAACTTTAGCCAGGGGCCAGTCCCGCAGTCGTGTAGGCTGGAGCTGCTTC |
| | | codA_LR_R1 | M | R | GAAATTACCGTTTGTAATCGATGGCTTCTGGCTGCTCCAGATATACGGTGCATATGAATATCCTCCTTAG |
| | | HcodB_F1 | S | F | CCCCCACCTTTTTGCACTCA |
| | | codA_R1 | S | R | CCACTACTGGAGAGAACGGC |
| pkd3 | | pKD3_C1 | S | F | TTATACGCAAGGCGACAAGG |
| | | pKD3_C2 | S | R | GATCTTCCGTCACAGGTAGG |
| pkd4 | | pKD4_K1 | S | F | TGTCCAGATAGCCCAGTAG |
| | | pKD4_K2 | S | R | CCTTCTATGAAAGGTTGGGC |
| lacZ | | lacZ_LR_F1 | M | F | ACAGCTATGACCATGATTACGGATTCACTGGCCGTCGTTTTACAACGTCGGTGTAGGCTGGAGCTGCTTC |
| | | lacZ_LR_R1 | M | R | ATTATTATTTTTGACACCAGACCAACTGGTAATGGTAGCGACCGGCGCTCCATATGAATATCCTCCTTAG |
| purA – pGEN-MCS | | purA_C_pGEN_F1 | C | F | TAAGCAggatccCATCCGTAGCCTGCGTGCTTA* |
| | | purA_C_pGEN_R1 | C | R | TGCTTAccatggttaCGCGTCGAACGGATCGCG* |
| purA – pCA24N | | purA_C_pCA24N_F1 | C | F | TAAGCAGGATCCatgGGTAACAACGTCGTCGTA |
| | | purA_C_pCA24N_R1 | C | R | TGCTTAGAGCTCttaCGCGTCGAACGGATCGCG |
| purA – pGEN-MCS | | purA_C_pGEN_Verify_F1 | C | F | GGCACTTGCTCACGCTCTG |
| | | purA_C_pGEN_Verify_R1 | C | R | GTGGTCACGCTTTTCGTTGG |
| purA – pCA24N | | purA_C_pCA24N_Verify_F1 | C | F | GATAACAATTTCACACAGAATTCATTAAAGAG |
| | | purA_C_pCA24N_Verify_R1 | C | R | CAAATCCAGATGGAGTTCTGAGG |
| purA | A4420 | purA_C_R | C | R | TTACCCTGCTTGCAGAGGAA |

*Lower case bases indicate restriction enzyme digest sites.

| Table S3 – Bacterial Strains | | |
|---|---|---|
| Strain | Description | Reference |
| E. coli | Commensal E. coli strain HS | (*1*) |
| Klebsiella pneumoniae | ATCC 43816 KPPR1 | (*2*) |
| Adherent Invasive E. coli (AIEC) | LF82, isolated from Crohn's Disease patient | (*3*) |
| Salmonella enterica serovar Typhimurium | WT Salmonella strain SL1344 | (*4*) |
| CJA116 | E. coli HS *ΔpurEK,* polar mutant, chloramphenicol resistant | This study |
| CJA117 | E. coli HS *ΔpurEK,* non-polar mutant | This study |
| SJB001 | E. coli HS *ΔpurA,* polar mutant, kanamycin resistant | This study |
| SJB014 | E. coli HS *ΔpurA,* non-polar mutant | This study |
| SJB003 | E. coli HS *ΔpurEK ΔpurA,* polar mutant, kanamycin resistant | This study |
| SJB017 | E. coli HS *ΔpurEK ΔpurA,* non-polar mutant | This study |
| CJA125 | E. coli HS *ΔpurMN,* polar mutant, chloramphenicol resistant | This study |
| CJA128 | E. coli HS *ΔpurMN,* non-polar mutant | This study |
| PC185 | E. coli HS *ΔpurEK ΔghxQ,* polar mutant, chloramphenicol resistant | This study |
| PC196 | E. coli HS *ΔpurEK ΔghxQ,* non-polar mutant | This study |
| PC171 | E. coli HS *ΔpurEK ΔghxP,* polar mutant, resistant | This study |
| PC177 | E. coli HS *ΔpurEK ΔghxP,* non-polar mutant | This study |
| PC194 | E. coli HS *ΔpurEK ΔghxP ΔghxQ,* polar mutant , chloramphenicol resistant | This study |
| PC200 | E. coli HS *ΔpurEK ΔghxP ΔghxQ,* non-polar mutant | This study |
| SJB023 | E. coli HS *Δapt,* polar mutant, kanamycin resistant | This study |
| SJB038 | E. coli HS *Δapt,* non-polar mutant | This study |
| SJB030 | E. coli HS *Δhpt,* polar mutant, kanamycin resistant | This study |
| SJB042 | E. coli HS *Δhpt,* non-polar mutant | This study |
| SR064 | E. coli HS *Δapt Δhpt,* polar mutant, kanamycin resistant | This study |
| SJB050 | E. coli HS *Δapt Δhpt,* non-polar mutant | This study |
| SJB025 | E. coli HS *ΔpurEK Δapt,* polar mutant, kanamycin resistant | This study |
| SJB040 | E. coli HS *ΔpurEK Δapt,* non-polar mutant | This study |
| SJB031 | E. coli HS *ΔpurEK Δhpt,* polar mutant, kanamycin resistant | This study |
| SJB044 | E. coli HS *ΔpurEK Δhpt,* non-polar mutant | This study |
| PC241 | E. coli HS *ΔpyrBI,* polar mutant, chloramphenicol resistant | This study |
| PC243 | E. coli HS *ΔpyrBI,* non-polar mutant | This study |
| PC255 | E. coli HS *ΔpyrF,* polar mutant, kanamycin resistant | This study |
| SJB052 | E. coli HS *ΔpyrF,* non-polar mutant | This study |
| SR005 | E. coli HS *ΔcodBA,* polar mutant, kanamycin resistant | This study |
| SR011 | E. coli HS *ΔcodBA,* non-polar mutant | This study |

| | | |
|---|---|---|
| PC247 | E. coli HS *ΔuraA,* polar mutant, chloramphenicol resistant | This study |
| PC251 | E. coli HS *ΔuraA,* non-polar mutant | This study |
| PC245 | E. coli HS *Δupp,* polar mutant, kanamycin resistant | This study |
| PC249 | E. coli HS *Δupp,* non-polar mutant | This study |
| PC048 | E. coli HS *ΔpurEK ΔpunC,* polar mutant, chlormaphenicol resistant | This study |
| PC056 | E. coli HS *ΔpurEK ΔpunC,* non-polar mutant | This study |
| PC093 | E. coli HS *ΔpurEK ΔpunC ΔadeP,* polar mutant, chloramphenicol resistant | This study |
| SR040 | E. coli HS *ΔpurEK ΔpunC ΔadeP,* non-polar mutant | This study |
| SR047 | E. coli HS *ΔpurEK ΔpunC ΔadeP ΔnupG,* polar mutant, chloramphenicol resistant | This study |
| SR052 | E. coli HS *ΔpurEK ΔpunC ΔadeP ΔnupG,* non-polar mutant | This study |
| PC122 | E. coli HS *ΔpurEK ΔpunC ΔadeP ΔnupG ΔghxP,* polar mutant, chloramphenicol resistant | This study |
| PC125 | E. coli HS *ΔpurEK ΔpunC ΔadeP ΔnupG ΔghxP,* non-polar mutant | This study |
| PC133 | E. coli HS *ΔpurEK ΔpunC ΔadeP ΔnupG ΔghxP ΔnupX,* polar mutant, chloramphenicol resistant | This study |
| PC136 | E. coli HS *ΔpurEK ΔpunC ΔadeP ΔnupG ΔghxP ΔnupX,* non-polar mutant | This study |
| PC141 | E. coli HS *ΔpurEK ΔpunC ΔadeP ΔnupG ΔghxP ΔnupX ΔghxQ,* polar mutant, chloramphenicol resistant | This study |
| PC146 | E. coli HS *ΔpurEK ΔpunC ΔadeP ΔnupG ΔghxP ΔnupX ΔghxQ,* non-polar mutant | This study |
| SR007 | E. coli HS *ΔlacZ,* polar mutant, chloramphenicol resistant | This study |
| PC055 | E. coli HS *ΔpurEK ΔlacZ,* polar mutant, kanamycin resistant | This study |
| PC118 | E. coli HS *ΔpurA ΔlacZ,* polar mutant, kanamycin resistant | This study |
| PC120 | E. coli HS *ΔpurEK ΔpurA ΔlacZ,* polar mutant, chloramphenicol resistant | This study |
| PC230 | E. coli HS *ΔghxP ΔlacZ,* polar mutant, kanamycin resistant | This study |
| PC169 | E. coli HS *Δhpt ΔlacZ,* polar mutant, kanamycin resistant | This study |
| PC175 | E. coli HS *Δhpt Δapt ΔlacZ,* polar mutant, kanamycin resistant | This study |
| PC234 | E. coli HS WT + pGEN-MCS empty vector control | This study |
| PC187 | E. coli HS *ΔpurA* (SJB014) + pPC021, *purA* insert into pGEN-MCS vector | This study |
| PC227 | E. coli HS *ΔpurA* (SJB014) + pGEN-MCS empty vector control | This study |
| PC179 | E. coli HS + pCA24N empty vector control | This study |
| PC192 | E. coli HS *ΔpurA* (SJB014) + pPC015, *purA* insert into pCA24N vector | This study |

| | | |
|---|---|---|
| PC184 | E. coli HS *ΔpurA* (SJB014) + pCA24N empty vector control | This study |


1. M. M. Levine, D. R. Nalin, R. B. Hornick, E. J. Bergquist, D. H. Waterman, C. R. Young, S. Sotman, B. Rowe, ESCHERICHIA COLI STRAINS THAT CAUSE DIARRHŒA BUT DO NOT PRODUCE HEAT-LABILE OR HEAT-STABLE ENTEROTOXINS AND ARE NON-INVASIVE. *The Lancet* **311**, 1119–1122 (1978).

2. I. A. J. M. Bakker-Woudenberg, J. C. van Den Berg, T. B. Vree, A. M. Baars, M. F. Michel, Relevance of serum protein binding of cefoxitin and cefazolin to their activities against Klebsiella pneumoniae pneumonia in rats. *Antimicrob. Agents Chemother.* **28**, 654–659 (1985).

3. A. Darfeuille-Michaud, C. Neut, N. Barnich, E. Lederman, P. Di Martino, P. Desreumaux, L. Gambiez, B. Joly, A. Cortot, J. F. Colombel, Presence of adherent Escherichia coli strains in ileal mucosa of patients with Crohn's disease. *Gastroenterology* **115**, 1405–1413 (1998).

4. S. K. Hoiseth, B. A. D. Stocker, Aromatic-dependent Salmonella typhimurium are non-virulent and effective as live vaccines. *Nature 1981 291:5812* **291**, 238–239 (1981).

| Table S4: Plasmids | | | |
| --- | --- | --- | --- |
| Plasmid | Description | Use | Reference |
| pKD46 | Lambda-red recombinase | Mutagenesis | (*1*) |
| pKD4 | Kanamycin resistant helper plasmid | Mutagenesis | (*1*) |
| pKD3 | Chloramphenicol resistant helper plasmid | Mutagenesis | (*1*) |
| pCP20 | Flippase | Mutagenesis | (*1*) |
| pGEN-MCS | Always on | Complementation | (*2*) |
| pAC24N | Arabinose inducible | Complementation | (*3*) |
| pPC021 | purA insert into pGEN-MCS | Complementation | This study |
| pPC015 | purA insert into PAC24N | Complementation | This study |


1. K. A. Datsenko, B. L. Wanner, One-step inactivation of chromosomal genes in Escherichia coli K-12 using PCR products. *Proc. Natl. Acad. Sci. U. S. A.* **97**, 6640–6645 (2000).

2. M. C. Lane, C. J. Alteri, S. N. Smith, H. L. T. Mobley, Expression of flagella is coincident with uropathogenic Escherichia coli ascension to the upper urinary tract. *Proc. Natl. Acad. Sci. U. S. A.* **104**, 16669–16674 (2007).

3. R. H. Wilson, S. K. Morton, H. Deiderick, M. L. Gerth, H. A. Paul, I. Gerber, A. Patel, A. D. Ellington, S. P. Hunicke-Smith, W. M. Patrick, Engineered DNA ligases with improved activities in vitro. *Protein Eng. Des. Sel.* **26**, 471–478 (2013).
